# SULTA4A1 modulates synaptic development and function by promoting the formation of PSD-95/NMDAR complex

**DOI:** 10.1101/583419

**Authors:** Lorenza Culotta, Benedetta Terragni, Ersilia Vinci, Alessandro Sessa, Vania Broccoli, Massimo Mantegazza, Chiara Verpelli

**Affiliations:** CNR Neuroscience Insitute, Milan, Milano; U.O. of Neurophysiopathology and Diagnostic Epileptology, Foundation Istituto di Ricerca e Cura a Carattere Scientifico (IRCCS) Neurological Institute Carlo Besta, Milan; Stem Cell and Neurogenesis Unit, Division of Neuroscience, San Raffaele Scientific Institute, 20132 Milan, Italy; Institute of Molecular and Cellular Pharmacology (IPMC), Laboratory of Excellence Ion Channel Science and Therapeutics (LabEx ICST), CNRS UMR7275 and University of Nice-Sophia Antipolis, Valbonne

**Author notes:** Corresponding author: Chiara Verpelli CNR Neuroscience Institute Via Vanvitelli 32 20129 Milano.

## Abstract

Sulfotransferase 4A1 (SULT4A1) is a cytosolic sulfotransferase, that is highly conserved across species and extensively expressed in the brain. However, the biological function of SULT4A1 is unclear. SULT4A1 has been implicated in several neuropsychiatric disorders, such as Phelan-McDermid Syndrome and schizophrenia. Here, we investigate the role of SULT4A1 within neuron development and function. Our data demonstrate that SULT4A1 modulates neuronal branching complexity and dendritic spines formation. Moreover, we show that SULT4A1, by negatively regulating the catalytic activity of Pin1 towards PSD-95, facilitates NMDAR synaptic expression and function. Finally, we demonstrate that the pharmacological inhibition of Pin1 reverses the pathological phenotypes of SULT4A1 knockdown neurons by specifically restoring dendritic spine density and rescuing NMDAR-mediated synaptic transmission. Together, these findings identify SULT4A1 as a novel player in neuron development and function by modulating dendritic morphology and synaptic activity.

## Introduction

Cytosolic Sulfotransferase 4A1 is member of the cytosolic sulfotransferases (SULT) superfamily, a class of enzymes that catalyze sulfonation reactions by transferring the sulfonate group from 3’-phosphoadenosine 5’-phosphosulfate (PAPS) to different endogenous and exogenous substrates(Falany et al., 2000, Negishi et al., 2001). SULT4A1 shows a remarkable degree of cross-species similarity, suggesting highly conserved biological function(Blanchard et al., 2004, Minchin et al., 2008). Of note, SULT4A1 has an atypical enzymatic domain structure, as revealed by a recent crystal structure(Allali-Hassani et al., 2007), that may affect PAPS binding and substrate specificity: in fact, no sulfonation activity has been detected with potential sulfate donors or substrates(Allali-Hassani et al., 2007, Falany et al., 2000), suggesting SULT4A1 may not exhibit classical catalytic activity *in vivo* or that the functional enzyme may be active as a component of a multi-enzyme complex(Falany et al., 2000). Thus, in absence of any known substrate, the biological function of SULT4A1 remains unclear.

SULT4A1 tissue distribution has been examined in both humans and rodents and has been demonstrated to be predominantly expressed within the brain, although limited expression of mRNA and protein is detectable in other organs, such as kidney, lung, liver and heart tissues(Alnouti et al., 2006, Sidharthan et al., 2014). In particular, the strongest protein expression has been detected in cerebral cortex, thalamus, cerebellum, and hippocampus(Liyou et al., 2003). In mouse, SULT4A1 mRNA expression is low in fetal brains and remains nearly unchanged until postnatal day 30, after which a marked increase in expression has been observed(Alnouti et al., 2006). Consistent with this, SULT4A1 protein expression was found to increase during neuronal differentiation(Idris et al., 2019). Notably, SULT4A1-KO mice display severe neurological defects including tremor, rigidity and seizure and the survival of pups during postnatal development is strongly affected by loss of SULT4A1(Garcia et al., 2018).

A growing body of evidence supports SULT4A1 as a key player in neuronal maturation, and that loss of SULT4A1 function can contribute towards neurodevelopmental disorders. For instance, SULT4A1 haploinsufficiency has been linked to neurological symptoms of patients with Phelan-McDermid Syndrome (PMS)(Disciglio et al., 2014), with SULT4A1 deletion strongly correlated to lower developmental quotient(Zwanenburg et al., 2016). In addition, single nucleotide polymorphisms (SNP) in SULT4A1 gene have been extensively linked to schizophrenia susceptibility as well as severity of both psychotic and intellectual impairment, and antipsychotic treatment response(Brennan and Condra, 2005, Meltzer et al., 2008, Ramsey et al., 2011). Altogether, these findings suggest a role of SULT4A1 in neuronal development and function and altered expression as a potential contributing factor in multiple neurodevelopmental disorders. Despite this body of supporting literature, the role of SULT4A1 within neuronal development and function remains unassessed. Our results provide evidence that SULT4A1 has an important role in the regulation of neuronal morphology and synaptic activity. Our data suggest that this effect is achieved through regulation, by direct protein-protein interaction, of the peptidyl-prolyl *cis-trans* isomerase Pin1 at a synaptic level. Pin1 is an enzyme that binds to phosphorylated serine/threonine-proline motifs and catalyzes *cis*-to-*trans* isomerization of prolines(Ranganathan et al., 1997, Shen et al., 1998), notably acting to regulate synaptic strength through regulation of post-synaptic scaffolds: for instance, in excitatory synapses Pin1 has been found to downregulate glutamatergic synaptic transmission by negatively modulating PSD-95/NMDAR complex formation(Antonelli et al., 2016). Here we demonstrate that, by sequestering Pin1, SULT4A1 facilitates the formation of the PSD-95/NMDAR complex in excitatory synapses, which is essential for NMDAR-mediated synaptic transmission and spines formation.

## Results

### Sulfotransferase 4A1 expression is increased during brain maturation

Considering that SULT4A1 has been implicated in neurodevelopmental and neuropsychiatric disorders such as schizophrenia(Brennan and Condra, 2005, Meltzer et al., 2008) and Phelan-McDermid Syndrome(Disciglio et al., 2014), a major point for this study was the evaluation of SULT4A1 neuronal expression during physiological maturation. To begin, SULT4A1 expression was analyzed during neuronal maturation in protein lysates harvested from rat cortical neurons in primary cultures at 1, 7, and 14 days in vitro (DIV1, 7 and 14). SULT4A1 expression was almost undetectable at DIV1 and increased during the neuronal maturation (Fig. 1a; DIV1: 0.03±0.01; DIV7: 0.18±0.02; DIV14: 0.38±0.03). Moreover, the immunostaining of cortical neurons revealed a localization of SULT4A1 to neuronal cell bodies and dendrites (Figure 1 figure supplement 1a). Then, SULT4A1 expression was evaluated *in vivo* during mouse brain development. Western blot (WB) analysis showed that SULT4A1 protein levels increased from post-natal day (P) 0 to P30. Moreover, high levels of SULT4A1 were maintained also in adult mice (P60), suggesting an important role of SULT4A1 both during brain maturation and adulthood (Fig. 1b; P0: 0.06; P7: 0.30; P14: 0.44; P21: 0.47; P30: 0.49; P60: 0.49).

**Figure 1.**
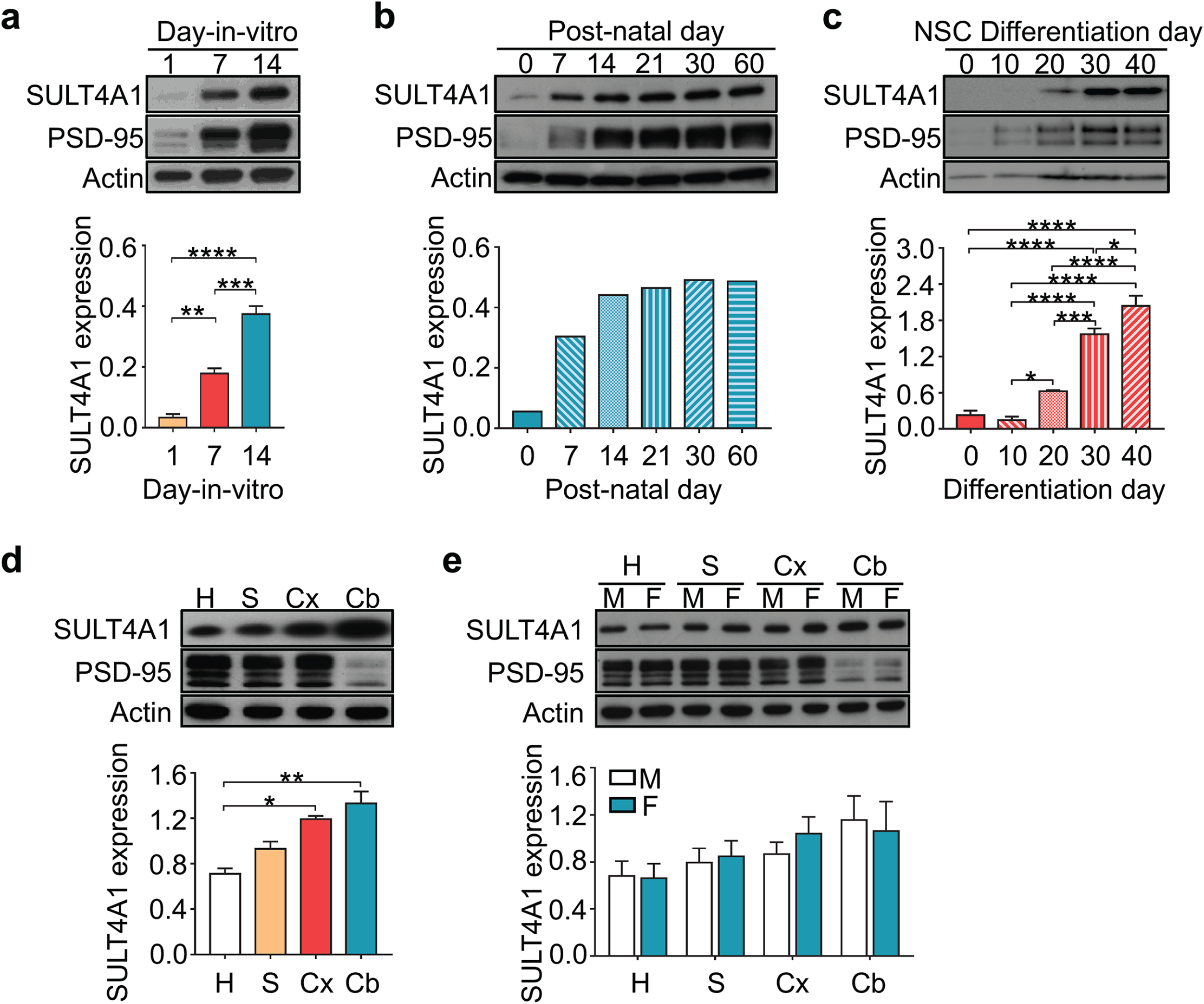
Physiological expression of SULT4A1 protein during development. (**a**) Representative western blots of total cell lysates derived from rat cortical neurons at different days-in-vitro (n=4 for all conditions; One-way ANOVA test, *P*<0.0001; Tukey’s *post-hoc* test, ** *P*<0.01, *** *P*<0.001, **** *P*<0.0001). (**b**) Biochemical analysis of total brain lysates derived from wild type male mice at different post-natal days (n=1 for all conditions). (**c**) Representative immunoblots of total cell lysates obtained from human neurons, sampled during differentiation from day 0 (NSC stage) until day 40 (mature neuron stage) (n=3 for all conditions; One-way ANOVA test, *P*<0.0001; Tukey’s *post-hoc* test, * *P*<0.05, *** *P*<0.001, **** *P*<0.0001). (d) Representative western blots of protein lysates from hippocampus (H), striatum (S), cerebral cortex (Cx) and cerebellum (Cb) derived from adult (P60) wild type mice (n=4 for all conditions; Kruskal-Wallis test, *P*<0.0001; Dunn’s *post-hoc* test, * *P*<0.05; ** *P*<0.01). Area-specific expression of SULT4A1 was also compared between adult male (M) and female (F) mice (n=3 for all conditions; Two-way ANOVA, P=0.8362; Sidak’s *post-hoc* test) (e). Data represent Mean ± SEM Detailed statistical information for represented data is shown in Supplementary file 1 The following figure supplement is available for figure 1: Figure supplement 1. Physiological expression of SULT4A1 in primary cortical neurons and iPSC-derived neurons.

To determine SULT4A1 protein expression in human neurons, human induced pluripotent stem cells (iPSCs) were differentiated into Nestin-positive Neural Stem Cells (NSCs) and subsequently into neurons (Figure 1 figure supplement 1b) and analyzed for SULT4A1 protein levels during differentiation. Human SULT4A1 expression was found to be increased during differentiation from NSCs (day 0) to mature neurons (day 40) (Fig. 1c; day 0: 0.23±0.08; day 10: 0.14±0.06; day 20: 0.63±0.02; day 30: 1.57±0.1; day 40: 2.04±0.17).

SULT4A1 is known to have particularly strong expression in the brain but to date, the current knowledge of SULT4A1 area-specific expression is restricted to human and rat tissues(Liyou et al., 2003). In order to assess the tissue distribution in adult mouse brain, WB analysis of total lysates of hippocampus, striatum, cerebral cortex and cerebellum was performed on 2-month-old male mice. SULT4A1 was found to be highly expressed in all analyzed areas (Fig. 1d; H: 0.71±0.05; S: 0.93±0.06; Cx: 1.19±0.03; Cb: 1.33±0.10). Finally, no significant difference in area-specific protein expression was observed between male and female animals (Fig. 1e; H: M 0.69±0.12, F 0.67±0.12; S: M 0.80±0.12, F 0.85±0.13; Cx: M 0.87±0.10, F 1.05±0.14; Cb: M 1.16±0.20, F 1.07±0.24).

### SULT4A1 silencing reduces neuronal arborization and dendritic spine density

It is widely acknowledged that schizophrenia and autism spectrum disorders are often correlated to deficiencies in dendrites architecture and spine dynamics(Hung et al., 2008, Glausier and Lewis, 2013, Jiang et al., 2013). In order to evaluate the impact of SULT4A1 on neuronal morphogenesis, a specific shRNA for SULT4A1 (shSULT4A1) was designed and validated in SULT4A1 transfected HEK cells (Figure 2 figure supplement 1).

To assess the role of SULT4A1 in early neuronal development, cortical neurons were transfected at DIV7 with shSULT4A1 or control scrambled shRNA (shCtrl) (Fig. 2a). Dendrite morphology of transfected neurons was then evaluated by Sholl analysis at DIV14 (Fig. 2b). SULT4A1 silencing resulted in a simplification of neuronal branching, indicated by the reduction of branching points (condition factor: *P*<0.0001; condition by distance interaction: *P*<0.0001), primary dendrites (shCtrl: 7.30±0.39, shSULT4A1: 4.70±0.43, SULT4A1r: 7.3±0.45) and secondary dendrites (shCtrl: 11.60±0.94, shSULT4A1: 6.26±0.51, SULT4A1r: 9.55±0.65) (Fig. 2b-d). To further confirm the role of SULT4A1 in modulating dendritic arborization *in vivo, in utero* electroporation was performed on wild type mouse cortical neurons with shSULT4A1 or shCtrl. Brain slices from 1-month-old electroporated animals were analyzed by confocal microscopy to measure dendritic complexity (Fig. 2i). Sholl analysis showed that, as *in vitro*, the number of branching points is decreased in cortical neurons following SULTA4A1 knockdown (Fig. 2l; condition factor: *P*=0.0112, condition by distance interaction: *P*=0.0629).

**Figure 2.**
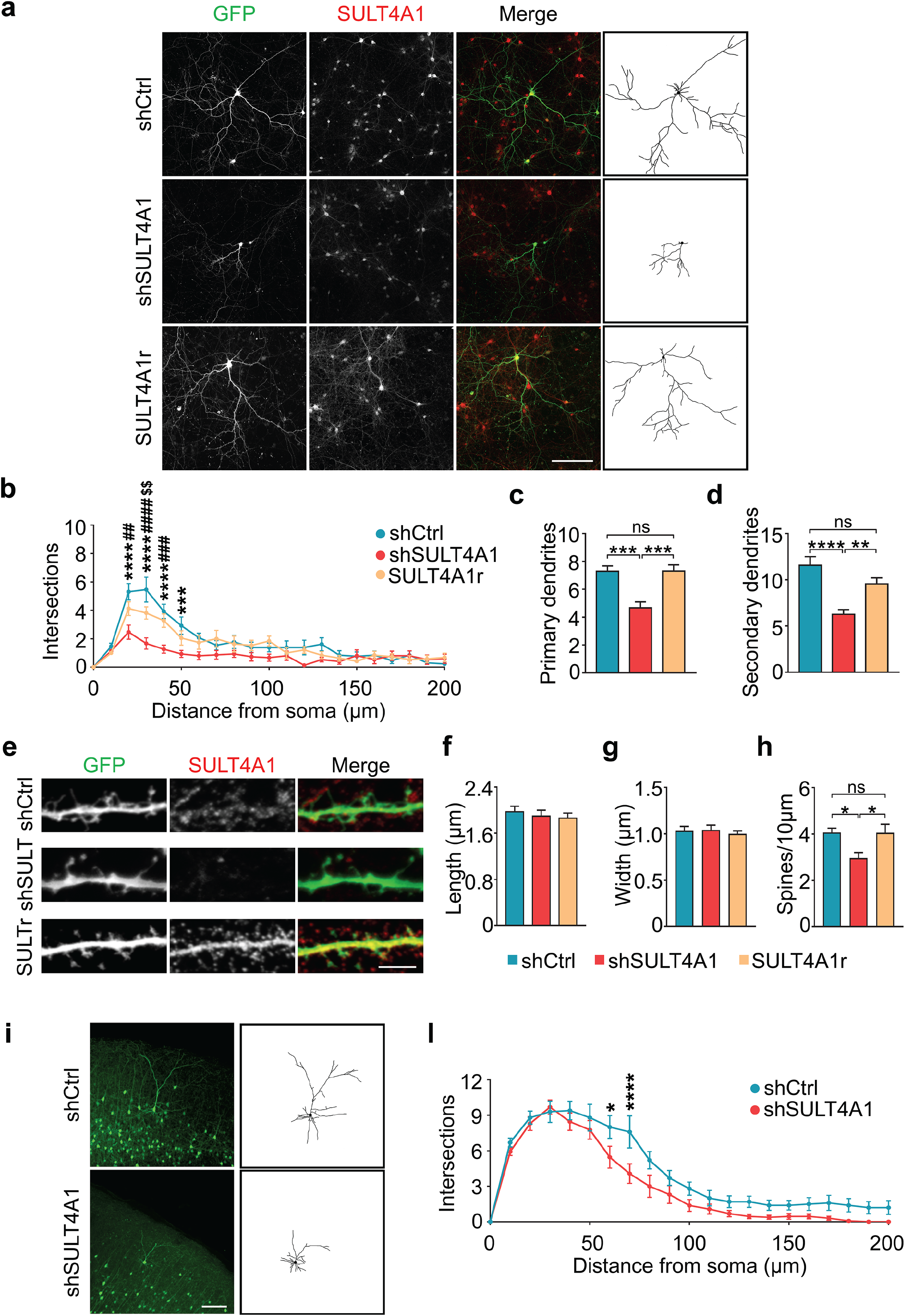
SULT4A1 silencing reduces neuronal branching and dendritic spine density. (**a**) Representative immunofluorescence images and relative traces of rat cortical neurons transfected with the scrambled shRNA (shCtrl), the SULT4A1-specific shRNA (shSULT4A1) or shSULT4A1 together with the resistant construct (SULT4A1r). Neurons were stained for SULT4A1 (red) to assess protein expression. Scale bar= 100 μm. (**b**) Results of Sholl analysis and quantification of the number of intersections, plotted against the distance from the soma (shCtrl n=13, shSULT4A1 n=15, SULT4A1r n=19; Two-way ANOVA, *P*<0.0001; Tukey’s *post-hoc* test, * shSULT4A1 vs shCtrl, *** *P*<0.001, **** *P*<0.0001; # shSULT4A1 vs SULT4A1r, ## *P*<0.01, ### *P*<0.001, #### *P*<0.0001: $ shCtrl vs SULT4A1r, $$ *P*<0.01). (**c-d**) Quantification of the number of primary (**c**) and secondary (**d**) dendrites (shCtrl n=21, shSULT4A1 n=18, SULT4A1r n=20; One-way ANOVA, *P*<0.0001, Tukey’s *post-hoc* test, ** *P*<0.01, *** *P*<0.001, **** *P*<0.0001, ns= not significant). (**e**) Representative immunofluorescence images showing dendritic spines of rat cortical neurons transfected with shCtrl, shSULT4A1 or shSULT4A1 together with the resistant plasmid. Neurons were stained for SULT4A1 (red) to assess protein expression. Scale bar= 10 μm. (**f-h**) Quantification of dendritic spines length (f), width (g) and number of spines per 10 μm of dendrite (h) (shCtrl n=10, shSULT4A1 n=10, SULT4A1r n=14; Kruskal-Wallis test, P=0.6092 for (f), P=0.8472 for (g), P=0.0081 for (h); Dunn’s *post-hoc* test, * *P*<0.05, ns= not significant). (**i**) Representative immunohistochemistry images from brain slices obtained from 30-day-old *in utero* electroporated mice, showing cortical neurons expressing shCtrl or shSULT4A1 and their relative traces. Scale bar= 100 μm. (**l**) Results of Sholl analysis and quantification of the number of intersections, plotted against the distance from the soma (shCtrl n=10, shSULT4A1 n=13; Two-way ANOVA, P=0.0629; Sidak’s *post-hoc* test, * *P*<0.05, **** *P*<0.0001). Data represent Mean ± SEM. Detailed statistical information for represented data is shown in Supplementary file 1 The following figure supplements are available for figure 2: Figure supplement 1. SULT4A1-specific shRNA validation. Figure supplement 2. SULT4A1 overexpression impairs dendritic arborization and spines.

The analysis of dendritic spine morphology revealed that SULT4A1-knocked down neurons exhibit a significant decrease in spines number (shCtrl: 4.07±0.18, shSULT4A1: 2.96±0.24, SULT4A1r: 4.05±0.36), but the morphology of the remaining spines was unchanged (f: shCtrl: 1.97±0.10, shSULT4A1: 1.90±0.10, SULT4A1r: 1.86±0.09; g: shCtrl: 1.03±0.05, shSULT4A1: 1.04±0.05, SULT4A1r: 1±0.03) (Fig. 2e-h). These morphological alterations were prevented by co-transfecting the shSULT4A1 with a shSULT4A1-resistant form of SULTA4A1 (SULT4A1r) (Figure 2 figure supplement 1), confirming the specificity of shSULTA4A1 knockdown. (Fig 2a-h).

To assess the role of SULTA4A1 dosage in modulating neuronal morphology, neurons were transfected at DIV7 with GFP or GFP plus SULT4A1 and stained at DIV14 with anti-SULT4A1 antibody to identify SULT4A1-overexpressing neurons (Figure 2 figure supplement 2a). SULT4A1 overexpression led to an overall redistribution of the branching points along the dendritic tree compared to wild type neurons: neurons overexpressing SULT4A1 presented fewer branching points close to the soma (<50 μm from soma) and an increased number of distal branching points (>100 μm from soma) (condition by distance interaction: *P*=0.0005) (Figure 2 figure supplement 2b), suggesting a reorganization of neuronal arborization. However, the number of primary (GFP: 7.7±0.36, Overexpression: 8.38±0.43) and secondary dendrites (GFP: 11.7±0.76, Overexpression: 13.05±0.86) was equal in the two conditions (Figure 2 figure supplement 2c-d). Interestingly, cortical neurons overexpressing SULT4A1 displayed a significant decrease in dendritic spine density (Figure 2 figure supplement 2h, spines/10μm: GFP 4.26±0.22, Overexpression 3.61±0.18), correlated to considerably longer spine morphology, as compared to wild type neurons, suggesting an immature phenotype (Figure 2 figure supplement. 2f, spine length, μm: GFP 1.95±0.08, Overexpression 2.71±0.19; Figure 2 figure supplement. 2g, spine width, μm: GFP 1±0.03, Overexpression: 1.02±0.03) (Figure 2 figure supplement. 2e-h).

These data demonstrate an important role for SULT4A1 in the regulation of both dendritic complexity and dendritic spine morphology.

### Functional modifications induced by SULT4A1 knockdown

Considering that SULT4A1 knockdown affected dendritic arborization and spine density, the effect of SULT4A1 silencing on the expression of relevant synaptic proteins was further investigated.

First, a lentiviral plasmid expressing shSULT4A1 was validated as effective in reducing the levels of endogenous SULT4A1 protein in cortical neurons (Fig. 3a; Ni: 1; shCtrl: 0.94±0.02, shSULT4A1: 0.08±0.02). Immunoblotting analysis of total lysates showed a significant decrease in NMDA receptor subunit GluN1 expression (shCtrl: 1, shSULT4A1: 0.81±0.06) and, simultaneously, a considerable increase of GAD65 protein level (shCtrl: 1, shSULT4A1: 1.38±0.16) in shSULT4A1-infected neurons (Fig. 3b).

**Figure 3.**
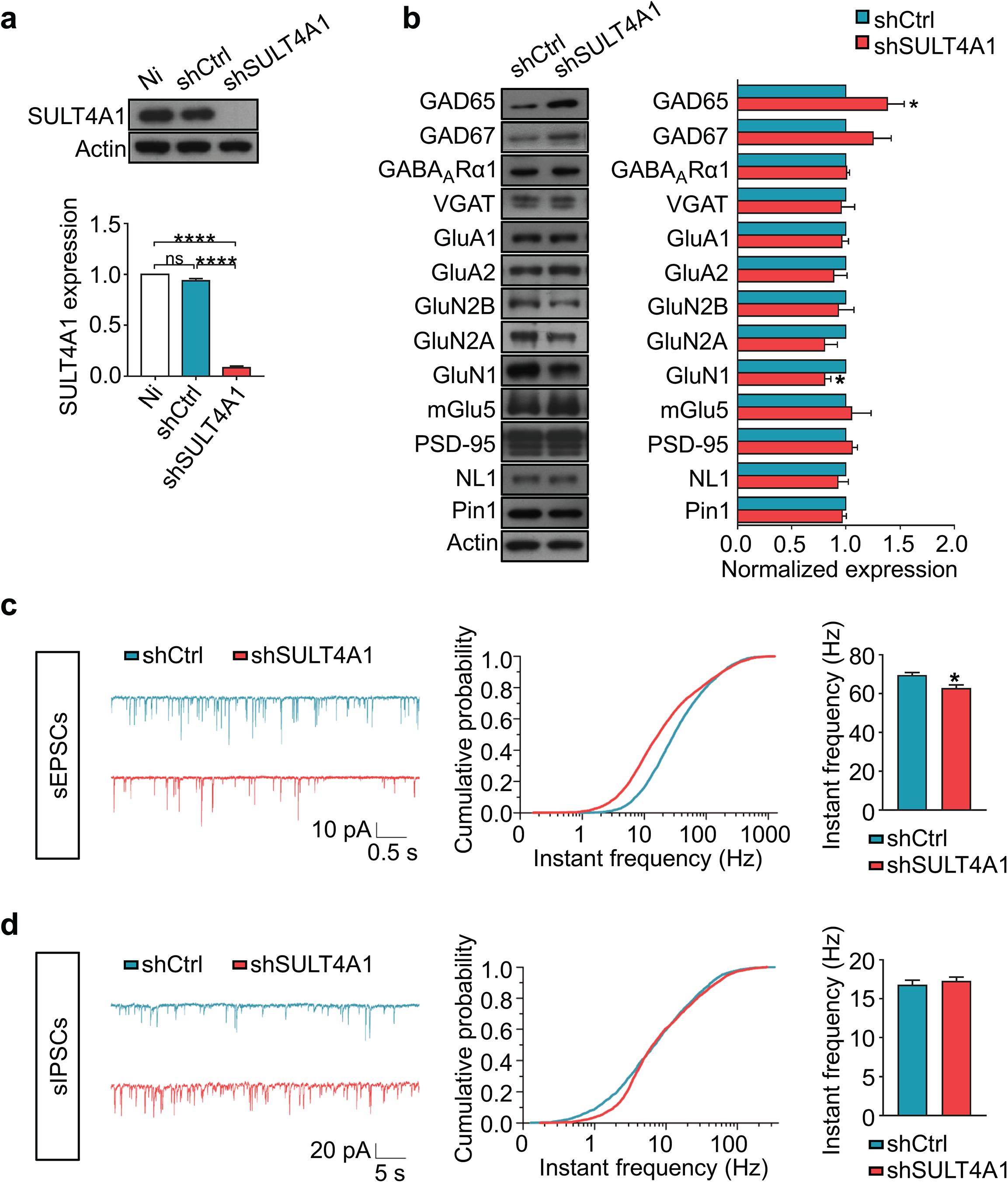
SULT4A1 knockdown alters synaptic transmission. (**a**) Representative immunoblots of total protein lysates derived from day-in-vitro 14 rat cortical neurons transduced with a lentivirus expressing shCtrl or shSULT4A1. Both conditions were compared with not infected condition (Ni) so to verify whether the infection itself could cause any alteration of SULT4A1 protein expression. SULT4A1 protein levels were normalized against the level of the not infected neurons (Ni n=7, shCtrl n=4, shSULT4A1 n=7; One-sample *t* test, **** *P*<0.0001, ns= not significant). (**b**) Representative western blots of total lysates from shCtrl- or shSULT4A1-transduced cortical neurons. Protein levels were normalized against the level of the shCtrl-transduced neurons. (shCtrl n≥3, shSULT4A1 n≥3; One-sample *t* test, * *P*<0.05). (**c**) Left: representative spontaneous excitatory postsynaptic currents (sEPSCs) recorded from shCtrl-(blue) or shSULT4A1-(red) transfected neurons. Center: plot of cumulative probability of currents frequency (shCtrl n=5, shSULT4A1 n=5; Kolmogorov-Smirnov test, *P*=4.16×10^−8^). Right: mean instantaneous frequencies plot (shCtrl n=5, shSULT4A1 n=5; Mann-Whitney test, * *P*<0.05). (**d**) Left: representative spontaneous inhibitory postsynaptic currents (sIPSCs) recorded from shCtrl-(blue) or shSULT4A1-(red) transfected neurons. Center: plot of cumulative probability of currents frequency (shCtrl n=5, shSULT4A1 n=6; Kolmogorov-Smirnov test, *P*=4.85×10^−8^). Right: mean instantaneous frequencies plot (shCtrl n=5, shSULT4A1 n=6; Mann-Whitney test). Data represent Mean ± SEM. Detailed statistical information for represented data (mean values, SEM, n, p) is shown in Supplementary file 1

To explore whether SULT4A1 silencing functionally affected glutamatergic and GABAergic activity, spontaneous post-synaptic currents were recorded from shCtrl or shSULT4A1 transfected neurons. SULT4A1 knockdown neurons displayed a significant reduction of spontaneous excitatory postsynaptic currents (sEPSC) frequency, reflected by the leftward shift of the cumulative probability curve and by the reduction of the median value (shCtrl: 69.6±1.3 Hz; shSULT4A1: 62.9±1.6 Hz) (Fig. 3c). Moreover, an increase of the frequency of spontaneous miniature inhibitory postsynaptic current (sIPSC) was observed in the 0.5-5 Hz range, reflected by the rightward shift of the cumulative probability curve, whereas the median value displayed a non-significant trend toward increased frequency (shCtrl: 16.8±0.6 Hz; shSULT4A1: 17.3±0.5 Hz) (Fig. 3d). These data suggest that SULT4A1 knockdown induced a decrease of glutamatergic synaptic activity and a moderate increase of GABAergic synaptic activity.

### SULT4A1 modulates NMDAR activity preventing Pin1-PSD-95 interaction

Our biochemical data indicate that the knockdown of SULT4A1 in neurons causes a significant decrease of GluN1. Therefore, total NMDAR-mediated currents (I_NMDA_) in neurons transfected with shSULT4A was assessed. A specific reduction of INMDA was observed, quantified as peak current density (shCtrl, pA/pF: 28.6±2; shSULT4A1: 22.3±2) (Fig. 4a). Immunoblot analysis revealed that the synaptosomal preparations derived from SULT4A1-deficient neurons exhibited significantly lower levels of GluN1 (shCtrl: 1; shSULT4A1: 0.83±0.02) and, albeit not statistically significant, a trend towards a reduction of GluN2B (p=0.1084, shCtrl: 1; shSULT4A1: 0.7852±0.10) and PSD-95 (p=0.2254, shCtrl 1; shSULT4A1: 0.93±0.05), (Fig. 4b).

**Figure 4.**
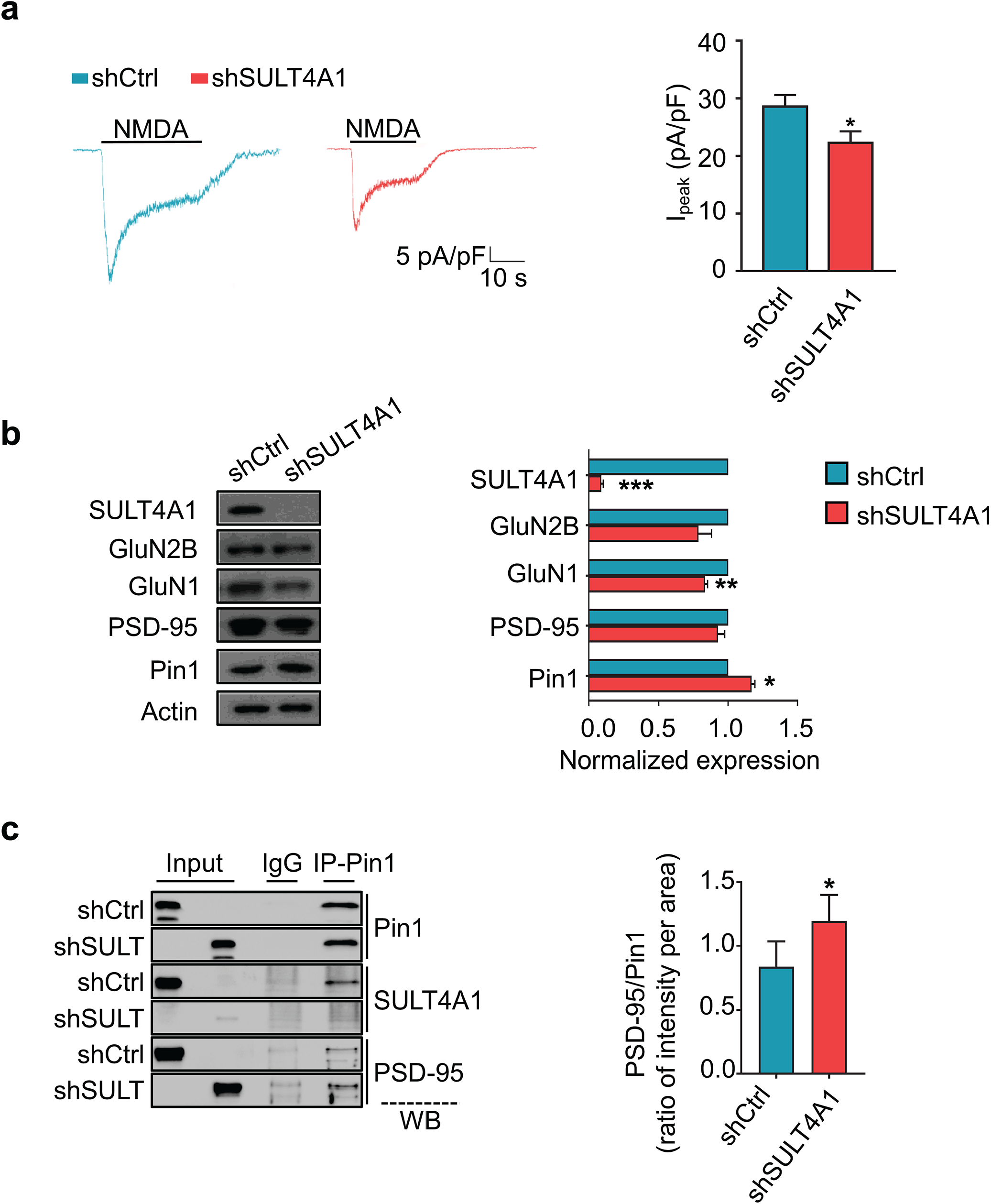
SULT4A1 interaction with Pin1 and its role in excitatory synapses. (**a**) Left: representative NMDA-mediated current (I_NMDA_) traces elicited by the perfusion of 100 μM NMDA from shCtrl-(blue) and shSULT4A1-(red) transfected neurons. Right: analysis of mean peak I_NMDA_ current densities in shCtrl- and shSULT4A1-transfected neurons (shCtrl n=9, shSULT4A1 n=13; Unpaired *t* test, * *P*<0.05). (**b**) Representative western blots of synaptosomal fractions obtained from cortical neurons transduced with shCtrl or shSULT4A1. Protein levels were normalized against the levels of the shCtrl-transduced neurons (shCtrl n≥3, shSULT4A1 n≥3; One-sample *t* test, * *P*<0.05, ** *P*<0.01, *** *P*<0.001). (**c**) Left: representative images of immunoprecipitation (IP) assay performed on synaptosomal fractions derived from neurons transduced with shCtrl or shSULT4A1. Proteins were precipitated using mouse anti-Pin1 or mouse IgG antibodies and nitrocellulose membranes were probed with anti-Pin1, anti-SULT4A1 and anti-PSD-95 antibodies (WB). Right: histogram showing the ratio between PSD-95 and Pin1 signals (PSD-95/Pin1) (shCtrl n=4, shSULT4A1 n=4; Unpaired *t* test, * *P*<0.05). Data represent Mean ± SEM. Detailed statistical information for represented data is shown in Supplementary file 1

Peptidyl-prolyl *cis-trans* isomerase Pin1 is highly expressed at excitatory synapses where it exerts a negative action on synaptic transmission by interfering with the PSD-95/GluN2B complex formation(Antonelli et al., 2016). Since it has been demonstrated that SULT4A1 is able to recruit Pin1 with the phosphoserine/threonine-proline motifs in its N-terminus(Mitchell and Minchin, 2009), the effects of SULT4A1 knockdown on synaptic expression of Pin1 was analyzed. A significant increase in Pin1 synaptic levels was evidenced by immunoblot analysis of synaptosomal preparations derived from SULT4A1-deficient neurons (shCtrl: 1; shSULT4A1: 1.17±0.02) (Fig. 4b).

To test if SULT4A1 directly affects the Pin1-PSD-95 interaction at synaptic levels, endogenous Pin1 was immunoprecipitated from synaptosomal preparations using an anti-Pin1 antibody, and the co-precipitated PSD-95 was visualized using an anti-PSD-95 antibody (Fig. 4c). With SULT4A1 silencing, the amount of PSD-95 co-precipitated by Pin1 was increased by 40% as compared to SULT4A1 expressing neurons (shCtrl: 0.83±0.10; shSULT4A1: 1.19±0.10). Thus, these results suggest that SULT4A1 is able to recruit Pin1 and, preventing its interaction with PSD-95, facilitates the formation of PSD-95/NMDAR complex. In conclusion, our data suggest that the reduction of NMDAR-mediated currents in SULT4A1-knocked down neurons is mediated by the increased interaction of Pin1 with PSD-95 in excitatory synapses.

### Morphological and functional rescue by PiB, a Pin1 inhibitor

As synaptic levels of Pin1 were found to be significantly increased following SULT4A1 knockdown, the role of Pin1 catalytic activity was assessed as a potential mechanism underlying altered glutamatergic transmission. The selective Pin1 inhibitor, PiB, was employed to inhibit Pin1 activity, in SULT4A1 knockdown neurons.

Cortical neurons transfected with shSULT4A1 or shCtrl at DIV7 were acutely treated with 2.5 μM PiB or vehicle (DMSO) at DIV12 for 48 hours. Acute treatment with PiB was able to restore the number of dendritic spines in SULT4A1-knocked down neurons to levels comparable to that of control neurons (shCtrl vehicle, spines/10μm: 5.67±0.30; shCtrl PiB: 6.12±0.42; shSULT4A1 vehicle: 4.33±0.20; shSULT4A1 PiB: 5.82±0.30) (Fig. 5a). To rescue dendritic arborization deficits, shSULT4A1 and shCtrl transfected neurons were treated every other day with 1 μM PiB or vehicle, starting at DIV7. However, chronic treatment with PiB had no significant effect on neuronal branching (shCtrl vehicle vs PiB: treatment factor P=0.4881, treatment by distance interaction P=0.9999; shSULT4A1 vehicle vs PiB: treatment factor P=0.3740, condition by distance interaction: P=0.9671) (Fig. 5b-c).

**Figure 5.**
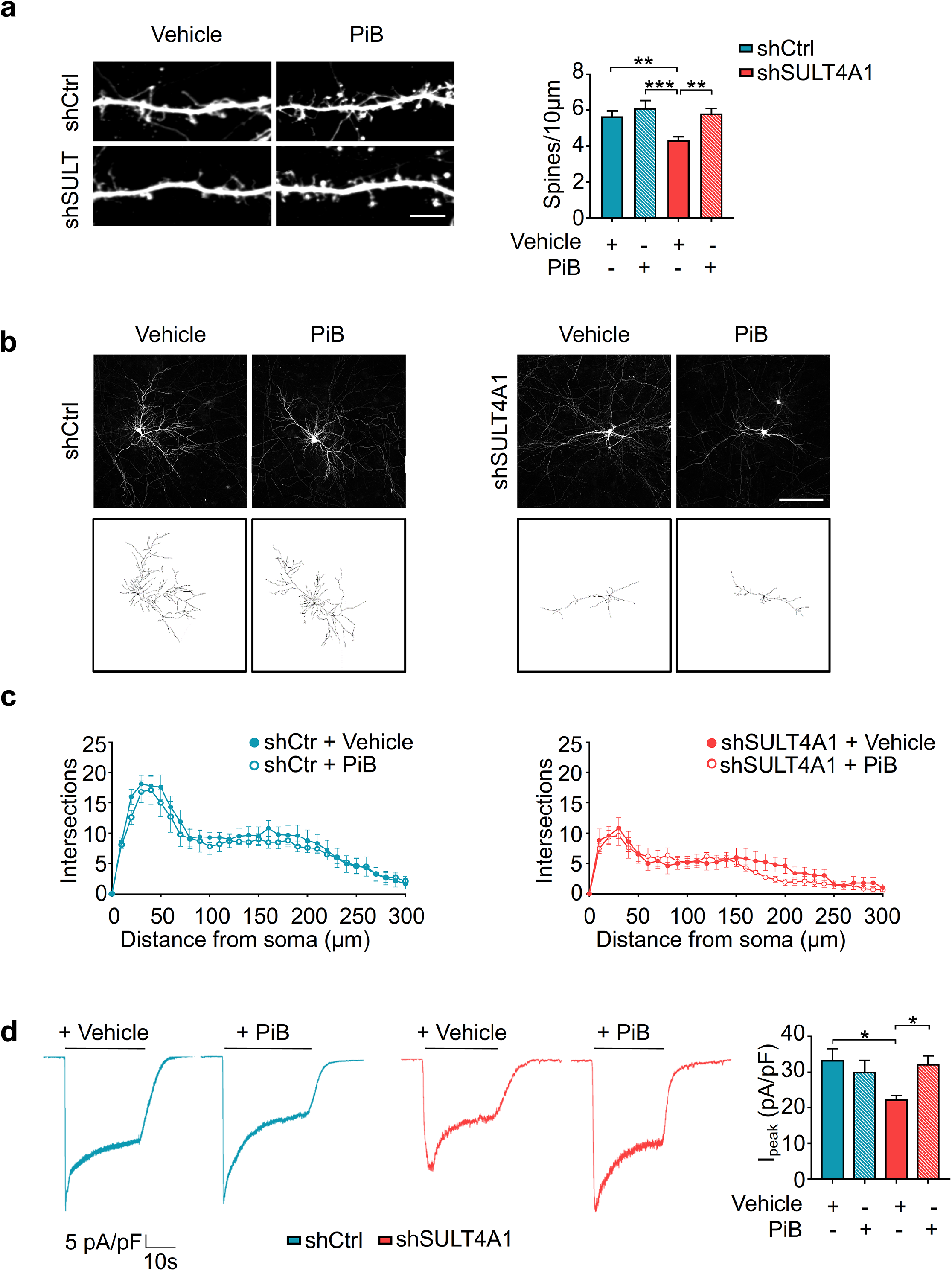
Partial rescue of shSULT4A1-dependent phenotype via Pin1 pharmacological inhibition. (**a**) Left: representative immunofluorescence images showing dendritic spines of rat cortical neurons transfected with shCtrl or shSULT4A1 and treated for 48h with 2.5 μM PiB, the pharmacological inhibitor of Pin1 catalytic activity, or vehicle (DMSO); scale bar=10μm. Right: quantification of the number of dendritic spines per 10 μm of dendrite (shCtrl + vehicle n=19, shCtrl + PiB n=16, shSULT4A1 + vehicle n=19, shSULT4A1 + PiB n=17; One-way ANOVA, *P*=0.0003; Tukey’s *post-hoc* test, ** *P*<0.01, *** *P*<0.001). (**b**) Representative immunofluorescence images and relative traces of rat cortical neurons transfected with shCtrl or shSULT4A1 and treated every other day with 1 μM PiB or vehicle, starting at DIV7. Scale bar= 100 μm. (**c**) Results of Sholl analysis and quantification of the number of intersections, plotted against the distance from the soma. shCtrl and shSULT4A1 conditions are displayed separately to clearly illustrate the effect of PiB treatment (Left: shCtrl + vehicle n=10, shCtrl + PiB n=11; Twoway RM ANOVA, *P*=0.9999; Sidak’s *post-hoc* test) (Right: shSULT4A1 + vehicle n=5, shSULT4A1 + PiB n=11; Two-way repeated measures ANOVA, *P*=0.9671; Sidak’s *post-hoc* test). (**d**) Left: representative NMDA-mediated current (I_NMDA_) traces from shCtrl-(blue) and shSULT4A1-(red) transfected neurons, treated for 48h with 2.5 μM PiB or vehicle. Right: analysis of mean peak I_NMDA_ current densities (shCtrl + vehicle n=12, shCtrl + PiB n=16, shSULT4A1 + vehicle n=6, shSULT4A1 + PiB n=16; One-way ANOVA, *P*=0.0082; Bonferroni’s *post-hoc* test, * *P*<0.05). Data represent Mean ± SEM. Detailed statistical information for represented data is shown in Supplementary file 1

Additionally, the effect of the acute PiB treatment on NMDAR-mediated currents was investigated. No significant effect was observed in neurons transfected with shCtrl. However, PiB treatment resulted in a 1.44-fold increase in peak current density in shSULT4A1-knocked down neurons (pA/pF, shCtrl vehicle: 33.1±3.3; shCtrl PiB: 29.8±3.5; shSULT4A1 vehicle: 22.2±1.2; shSULT4A1 PiB: 32±2.6) (Fig. 5d). Thus, Pin1 inhibition can fully rescue NMDA currents alteration induced by SULT4A1 loss of function.

These results suggest that SULT4A1, through direct interaction with Pin1 in dendritic spines, plays a novel role in regulating dendritic spine maturation and synaptic transmission.

## Discussion

SULT4A1 is a cytosolic sulfotransferase predominantly expressed in the brain and emerging as a genetic factor in a variety of neurodevelopmental diseases. However, the functional role of SULT4A1 within neuronal development is largely unknown since no known substrate or biological functions have been identified yet. SULT4A1 mRNA and protein expression were found to increase during mouse neurons and brain development(Alnouti et al., 2006, Hashiguchi et al., 2018, Idris et al., 2019), supporting a role for SULT4A1 in neuronal maturation. We have confirmed this timeline in murine models and demonstrate a similar increase in a human iPSC-derived neuronal model (Fig. 1a-c). Notably, we found that SULT4A1 is not only express in cytosol, mitochondrial and microsomal fractions(Garcia et al., 2018) but is also highly expressed in synaptosomal preparation, obtained from mouse cortical neurons (Fig. 4b).

SULT4A1-KO mice present with a severe and progressive neurologic phenotype, including tremor, absence seizures and ataxia, resulting in postnatal death(Garcia et al., 2018). In humans, polymorphisms in the SULT4A1 gene have been associated with susceptibility to schizophrenia(Brennan and Condra, 2005) and several intronic polymorphism were found in patients with worse cognitive performance(Meltzer et al., 2008). Interestingly, these SULT4A1 polymorphisms are believed to lead to a reduction of mRNA translatability(Brennan and Condra, 2005). Phelan-McDermid Syndrome (PMS) is a neurological disorder characterized by global developmental delay and autistic like behavior(Phelan and McDermid, 2012), due to deletions of the distal long arm of chromosome 22. Although it is widely recognized that deletion of SHANK3 gene, encoding a scaffold protein of the post-synaptic density(Naisbitt et al., 1999), is the main cause of the PMS neurological phenotypes(Durand et al., 2007), the wide clinical heterogeneity among PMS patients suggests that the haploinsufficiency of other genes in the 22q13 region, beside SHANK3, might contribute to cognitive and speech deficits associated with PMS. Approximately 30% of patients with PMS have a deletion encompassing SULT4A1(Sarasua et al., 2014) and infants with SULT4A1 deletion displayed a developmental quotient lower then patients showing two intact SULT4A1 alleles(Zwanenburg et al., 2016). Moreover, Disciglio et al. proposed SULT4A1 as a gene related to neurological symptoms of PMS patients(Disciglio et al., 2014). All these evidences and the high evolutionary conserved sequence in vertebrate brains(Allali-Hassani et al., 2007), strengthen the hypothesis that SULT4A1 plays an important role in central nervous system development and function.

Garcia et al. showed that SULT4A1 is expressed in the neurite projections of primary cortical neurons(Garcia et al., 2018). Our morphological data show that, in cultured neurons and *in vivo*, SULT4A1 modulates neuronal branching complexity and dendritic spine density, suggesting a role of SULT4A1 in neuronal maturation and synaptic plasticity. Deficits in dendritic arborization and spine density have been characterized in PMS and schizophrenia(Verpelli et al., 2011, Peca et al., 2011, Gouder et al., 2019, Glausier and Lewis, 2013, Russell et al., 2018). Interestingly, we found alteration in dendritic arborization and spine density both when we silence or overexpress SULT4A1 (Fig. 2 and Supplementary Fig. 3). These observations support expression of SULT4A1 at the appropriate level as crucial to ensure proper neuronal development and function. Thus, deficits in SULT4A1 expression, and the resulting decrease in arborization and spine density, may confer developmental deficits that contribute to the presentation or severity of PMS and schizophrenia.

Electrophysiological recordings in primary neurons showed that SULT4A1 modulates NMDAR-mediated synaptic transmission. SULT4A1 silencing induced a decrease in NMDA receptor subunit GluN1 expression (Fig. 3 and Fig. 4b) that was associated with a significant reduction in NMDA current amplitude (Fig. 4a).

The peptidyl-prolyl *cis-trans* isomerase Pin1, highly expressed in neuronal cells, has been identified as an interactor of SULT4A1(Mitchell and Minchin, 2009, Smet et al., 2005). However, the physiological role of this interaction is still unknown. Pin1 is a key regulator of synaptic plasticity(Antonelli et al., 2016), playing a central role in the prolyl isomerization of PSD-95. This function has been shown to negatively affect the formation of the PSD-95/NMDAR complex that is essential for targeting NMDARs to synapses(Antonelli et al., 2016). This role is supported by the observation that neurons derived from Pin1^−/−^ mice exhibit increased spine density and synaptic GluN1 and GluN2B content(Antonelli et al., 2016). In the absence of SULT4A1, we found a higher amount of Pin1 at synaptic sites associated with lower levels of GluN1 and GluN2B (Fig. 4b). Considering the known catalytic activity of Pin1 towards PSD-95(Antonelli et al., 2016), it is compelling to note that in co-immunoprecipitation experiments from cortical neurons knocked down for SULT4A1 we consistently detected an enhanced Pin1/PSD-95 complex formation (Fig. 4c). These results suggest that PSD-95-targeted prolyl isomerization mediated by Pin1 may be enhanced by SULT4A1 knockdown.

Notably, reduced NMDAR-mediated synaptic transmission caused by the absence of SULT4A1 expression can be rescued through pharmacological inhibition of Pin1 catalytic activity (Fig. 5d). In line with these results, Pin1 inhibition induces a significant rescue of dendritic spines number (Fig. 5a) and NMDA current (Fig. 5d) in SULT4A1 knockdown neurons. These results support a role for SULT4A1 in the regulation of PSD-95-targeted Pin1 activity, which in turn controls GluN1 synaptic levels and the formation or stability of dendritic spines. As NMDAR function is intricately linked to synaptic maturation and activity, it is compelling to hypothesize a direct link between GluN1 synaptic levels, dendritic spine density and the well characterized behavioral deficits within SULT4A1 knockout models(Garcia et al., 2018, Crittenden et al., 2015).

In conclusion, the present study reveals a novel role for SULT4A1 in modulating neuronal morphology and synaptic activity. We propose a model in which SULT4A1 acts to sequester Pin1 from the synapse, preventing its isomerase activity towards PSD-95. As PSD-95 prolyl-isomerization has been characterized to negatively modulate the synaptic content of NMDA receptors(Antonelli et al., 2016), this sequestration facilitates the formation of PSD-95/NMDAR complexes within dendritic spines, resulting in increased NMDAR-mediated synaptic activity and dendritic spine maturation (Fig. 6).

**Figure 6.**
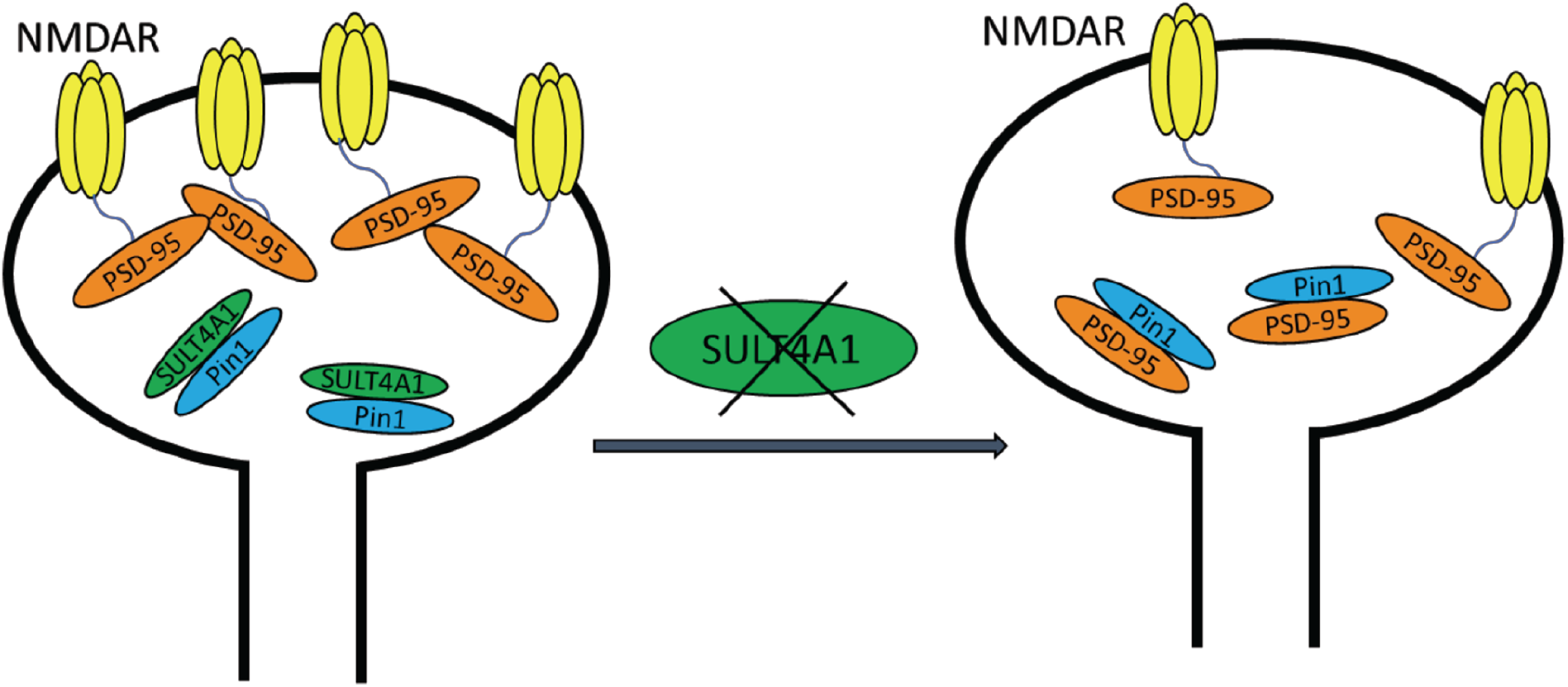
SULT4A1 modulates NMDAR synaptic expression and function. SULT4A1 interacts with Pin1 promoting the formation of the PSD-95/NMDAR complex in excitatory synapses. In absence of SULT4A1, Pin1 is free to interact with PSD-95 reducing NMDAR expression in synapses.

## Materials and Methods

### Constructs and virus generation

For RNA interference, a siRNA sequence targeting the SULT4A1 C-terminus was designed following GenScript siRNA Target Finder instructions (GenScript). Nucleotide sequence: AAGTGTGACCTCACGTTTGAC. The sequence was used to generate a short hairpin RNA (indicated as shSULT4A1 or shSULT) which was cloned into the pLVTHM-GFP vector(Wiznerowicz and Trono, 2003) using EcoRI and ClaI restriction sites. A scrambled form of shRNA was cloned into pLVTHM-GFP so to generate the control shRNA (indicated as shCtrl)(Heise et al., 2017). Previously described pFlag-SULT4A1(Mitchell and Minchin, 2009) was used for the overexpression experiments alongside pLVTHM-GFP vector as an overexpression control. Site directed mutagenesis was performed using QuikChange Lightning Site-Directed Mutagenesis Kit (Agilent Technologies) to generate a construct resistant to interference by shSULT4A1 (indicated as SULT4A1r or SULTr). In this construct three nucleotides (G825A, T828C, C831T) of the shSULT4A1 target site were altered, without changing the amino acid sequence of the resultant protein.

For viral transduction, genetically modified lentiviruses were produced as previously described(Lois et al., 2002, Naldini et al., 1996) and the production was carried out with 2^nd^ and 3^rd^ generation lentiviral transfer vectors.

### Animals

To prepare primary neuronal rat cultures, pregnant female rats (Rattus norvegicus) of the phylum Wistar strain were purchased from Charles River (Charles River Laboratories, Calco, Italy). C57BL/6 wild-type mice were purchased from Charles River. Mice and rats were housed under constant temperature (22 ± 1°C) and humidity (50%) conditions with a 12 hour light/dark cycle and were provided with food and water *ad libitum*. All experiments involving animals followed protocols in accordance with the guidelines established by the European Communities Council and the Italian Ministry of Health (Rome, Italy).

### Cell lines and transfections

HEK293T cells were cultured at 37°C and 5% CO_2_ atmosphere in DMEM (ThermoFisher) supplemented with fetal bovine serum (10%, ThermoFisher), L-Glutamine (2 mM, Euroclone), PenStrep (1%, ThermoFisher). Cells were transfected using Lipofectamine 2000 (ThermoFisher) and collected 24–48 hours after transfection.

### Primary neuronal cell culture

Low density rat cortical neuronal cultures were prepared from embryonic day (E) 18 rat embryos (Charles River) as previously described(Verpelli et al., 2010). Neurons were plated at 150-200 cells/mm^2^ density on 6- or 12-well plates (Euroclone) coated with 0.01 mg/ml poly-L-Lysine (Sigma-Aldrich). Neurons were cultured in Neurobasal (ThermoFisher) supplemented with the previously described B27(Chen et al., 2008). Cells were cultured on 6- and 12-well plates for protein biochemical analysis, whereas 12-well plates with acid-treated coverslips (VWR) were used for immunocytochemical or electrophysiological analysis. At day-in-vitro 7 (DIV7), neurons were lentivirally transduced or transfected using Lipofectamine 2000 (ThermoFisher). Biochemical, electrophysiological and morphological experiments were performed at DIV14.

#### PiB treatment

In order to inhibit Pin1 catalytic activity, cortical neurons were treated with PiB (diethyl-1,3,6,8-tetrahydro-1,3,6,8-tetraoxobenzol-phenanthroline-2,7-diacetate). PiB (Calbiochem) was resuspended in DMSO. As acute treatment, neurons were treated for 48h with 2.5 μM PiB and analyzed at DIV14; as chronic treatment, neurons were treated with 1 μM PiB every other day, from DIV7 to DIV14.

### Derivation of human iPSCs and neuronal differentiation

Human blood samples were collected according to a clinical protocol approved by the local Bioethical Committees of different medical centers. Participating individuals have been informed of the objectives of the study and signed an informed consent before inclusion in the study. Peripheral blood mononuclear cells (PBMCs) were isolated using Ficoll and growth in StemPro-34 SFM Medium (ThermoFisher), supplemented with L-Glutamine (2 mM, Euroclone), PenStrep (1%, ThermoFisher), SCF (100 ng/mL, ThermoFisher), FLT-3 (100 ng/mL, ThermoFisher), IL-3 (20 ng/mL, ThermoFisher), IL-6 (20 ng/mL, ThermoFisher). To generate human induced pluripotent stem cells (iPSCs), PBMCs were transduced with 2.0 Sendai virus particles containing four Yamanaka factors using the CytoTune-iPS Sendai Reprogramming Kit (ThermoFisher). After seven days, transduced cells were plated on cultures dishes coated with hESC-qualified matrigel (Corning) and grown in feeder-free conditions with Essential 8 medium (ThermoFisher). Three to four weeks after transduction, iPSCs colonies were manually picked for further expansion or analysis.

For neural stem cells (NSCs) derivation, iPSCs were detached with UltraPure EDTA (ThermoFisher) and plated on matrigel-coated 6-well plates in Essential 8 medium. After 24 hours, the spent medium was replaced with PSC Neural Induction Medium (ThermoFisher) and subsequently changed every other day following manufacturer’s instructions. On day 7 of neuronal induction, NSCs (P0) were harvested with StemPro Accutase (ThermoFisher) and plated on matrigel-coated plates for further expansion.

To obtain terminally differentiated neurons, proliferating NSCs were plated on matrigel-coated 6- or 12-well plates and cultured in Neurobasal medium supplemented with B27 without vitamin A (2%, ThermoFisher), PenStrep (1%, ThermoFisher), Glutamax (2mM, ThermoFisher), NT-3 (10ng/mL, Miltenyi Biotec), BDNF (10ng/mL, Miltenyi Biotec), GDNF (10ng/ml, Miltenyi Biotec), Retinoic Acid (1μM, Sigma-Aldrich) and growth for at least 40 days(Borroni et al., 2017). Half medium changes were performed every 2–3 days thereafter.

### Biochemical analysis

Cells or brain were lysed in pre-chilled buffered sucrose [0.32 M sucrose (Sigma-Aldrich)/ 4 mM HEPES-NaOH buffer (Sigma-Aldrich), pH 7.3, protease inhibitors (Sigma-Aldrich), phosphatase inhibitors (Roche)] and analyzed via Bradford protein assay (Bio-Rad) to assess protein concentration. For total lysate, proteins were directly solubilized in 4X loading buffer [(250 mM Tris, 40% glycerol, 0.008% bromophenol blue (all Sigma-Aldrich)]. In other cases, fractionation took place prior to solubilization to obtain a synaptosome-enriched fraction (P2), as previously published(Vicidomini et al., 2017).

For immunoblotting, primary antibodies and HRP-conjugated secondary antibodies were applied o/n and for 1 hour, respectively. Chemiluminescence was induced using an ECL kit (GE Healthcare) and densitometry was performed using Fiji/ImageJ (US National Institutes of Health).

### Immunoprecipitation assay

To evaluate the interaction between Pin1, SULT4A1 and PSD-95, primary neurons were lysed in NP-40 lysis buffer [1% NP-40, 50mM Tris pH 8.0, 150mM NaCl and protease inhibitor cocktail (all Sigma-Aldrich)] and subjected to synaptosomal fractionation as previously described(Vicidomini et al., 2017). P2 fractions were pre-cleared in NP-40 buffer for 4 hours and then incubated o/n at 4 °C with protein A-Sepharose beads (GE Healthcare, Milan, Italy) conjugated to 10 μg/ml of anti-Pin1 antibody (Santa Cruz Biotechnology) or matched control IgG (Sigma-Aldrich). The beads were then washed with 0.5% NP-40 buffer, re-suspended in loading buffer, warmed at 65 °C for 10 minutes and analyzed using SDS-PAGE. Samples of pre-cleared P2 fractions were collected before immunoprecipitation and used as input signals.

### Electrophysiological recordings

Whole-cell patch-clamp recordings in voltage-clamp configuration were performed at room temperature (~25°C) on DIV14 neurons using a Multiclamp 700A amplifier and pClamp 10.5 software (Molecular Devices, Sunnyvale, CA, USA). Signals were filtered at 1 kHz and sampled at 10 kHz. Postsynaptic currents were recorded at −70 mV in an external bath solution containing (mM): 129 NaCl, 1.25 NaH_2_PO_4_, 35 glucose, 1.8 MgSO_4_, 1.6 CaCl_2_, 3 KCl and 10 HEPES, pH 7.4 with NaOH. Spontaneous excitatory postsynaptic currents (sEPSCs) were recorded in the presence of gabazine (125 μM) with an intracellular pipette solution containing (mM): 120 K-gluconate, 15 KCl, 2 MgCl_2_, 0.2EGTA, 10HEPES, 20 phosphocreatine-tris, 2 ATP-Na_2_, 0.2 mM GTP-Na_2_, 0.1 leupeptin and 3 lidocaine N-ethyl bromide, pH 7.2 with KOH. Spontaneous inhibitory postsynaptic currents (sIPSCs) were recorded in the presence of kynurenic acid (3 mM) with an intracellular pipette solution containing (mM): 135 CsCl, 2 MgCl_2_, 0.2 EGTA, 10 HEPES, 20 phosphocreatine-tris, 2 ATP-Na_2_, 0.2 mM GTP-Na_2_ and 0.1 leupeptin, 3 lidocaine N-ethyl bromide, pH 7.2 with KOH; in these conditions, the Cl^−^ reversal potential was 0 mV, thus allowing to record at hyperpolarized potentials (−70 mV). Signals were filtered at 1 kHz and sampled at 10 kHz. The quantification of the instantaneous frequency of postsynaptic currents was performed on 2 min-long recordings with Clampfit 10.5.

NMDA-mediated currents (I_NMDA_) were elicited by the perfusion of 100 μM NMDA and recorded at −70 mV with an external bath solution containing (mM): 150 NaCl, 3 KCl, 3 CaCl_2_, 10 HEPES, 8 glucose, 0.1 D-Serine, 0.001 tetrodotoxin, 0.01 DNQX, 0.01 picrotoxin and 0.001 CGP 55845, pH 7.4 with NaOH, and an internal pipette solution containing (mM): 135 CH_3_O_3_SCs, 8 NaCl, 0.5 EGTA, 10 HEPES, 0.3 GTP-Na_2_ and 4 ATP-Mg, pH 7.2 with CsOH. NMDA application was performed with a local fast perfusion system controlled by electronic valves (RSC-200 Rapid Solution Changer, Bio-Logic Science Instruments); the tip of a multichannel manifold was placed at a distance of 0.2 mm from the patched neuron, allowing a fast solution exchange. To avoid drug accumulation, 10 minutes of wash out was applied after each recording. Peak I_NMDA_ were normalized to the membrane capacitance to obtain the current densities for statistical analysis.

### *In utero* electroporation, section preparation and immunohistochemistry

*In utero* electroporation was employed to deliver shCtrl- and shSULT4A1-expressing vectors to the ventricular RGCs of CD1 mouse embryos as previously described(Sessa et al., 2008). Briefly, uterine horns of E 13.5 pregnant dams were exposed by midline laparotomy after anesthetization with Avertin (312 mg/kg, Sigma-Aldrich). 1 μl of DNA plasmid(s) corresponding to 3 μg mixed with 0.03% fast-green dye in PBS was injected in the telencephalic vesicle using a pulled micropipette through the uterine wall and amniotic sac. 7 mm platinum tweezer-style electrodes were placed outside the uterus over the telencephalon and 5 pulses of 40 V, 50 ms in length, were applied at 950 ms intervals by using a BTX square wave electroporator. The uterus was then replaced in the abdomen, the cavity was filled with warm sterile PBS and the abdominal muscle and skin incisions were closed with silk sutures.

At post-natal day 30, GFP-positive brains were dissected for subsequent analyses. To this purpose, animals were anesthetized with an intraperitoneal injection of Avertin and perfused with 5% sucrose and 4% paraformaldehyde. Then, brains were left overnight in 4% paraformaldehyde, followed by incubation in 30% sucrose. Finally, brains were included in cryomolds with Tissue-Tek OCT compound (Sakura) and stored at −80°C until cryostat sectioning. 50 μm-thick slices were collected on polysine microscope adhesion slides (ThermoFisher) and then incubated in blocking solution (3% BSA, 10% goat serum, 0.4% Triton-X-100, diluted in PBS). Primary and fluorophore-conjugated secondary antibodies were diluted in blocking solution and applied respectively o/n at 4°C and 1 hour at room temperature. Glass coverslips were mounted on slides with Mounting Medium (VectaShield).

All procedures were approved by the Italian Ministry of Health and the San Raffaele Scientific Institute Animal Care and Use Committee in accordance with the relevant guidelines and regulations.

### Immunocytochemistry

Cells were fixed in 4% paraformaldehyde, 4% sucrose in PBS [136.8 mM NaCl, 2.68 mM KCl, 10.1 mM Na_2_HPO_4_ and 1.76 mM KH_2_PO_4_, pH 7.4 (all Sigma-Aldrich)] at room temperature for 10 minutes. Primary antibodies were diluted in homemade gelatin detergent buffer (GDB) [30 mM phosphate buffer, pH 7.4, 0.2% gelatin, 0.5% Triton X-100, 0.8 M NaCl (all Sigma-Aldrich)] and applied o/n at 4°C. Secondary antibodies conjugate with fluorophores (Jackson ImmunoResearch Laboratories) were also diluted in GDB buffer and applied for 1h. DAPI staining (ThermoFisher) was carried out at a final concentration of 0.5 μg/ml. Coverslips were mounted on pre-cleaned microscope slides using Mowiol mounting medium(Osborn and Weber, 1982).

### Image acquisition and processing

Confocal images were obtained using LSM 510 Meta confocal microscope (Carl Zeiss) with Zeiss 63X, 40X or 20X objectives at a resolution of 1024×1024 pixels. Images represent averaged intensity Z-series projections of 2-7 individual images taken at depth intervals of around 0.45 μm. For dendritic spine analysis, morphometric measurements were performed using Fiji/ImageJ software (US National Institutes of Health). Individual dendrites were selected randomly, and their spines were traced manually. Maximum length and head width of each spine were measured and archived automatically.

For neuronal arborization analysis, primary and secondary dendrites were measured manually while Sholl analysis was performed using Fiji/ImageJ software. Branching points were counted and plotted against distance from the soma.

### Antibodies

The following primary antibodies were used: mouse anti-β-actin (Sigma-Aldrich, A5316), mouse anti-βIII-tubulin (Sigma-Aldrich, MAB1637), mouse anti-GABA_A_Rα1 (NeuroMab, 75-136), mouse anti-GAD65 (Synaptic System, 198 111), mouse anti-GAD67 (Santa Cruz Biotechnology, sc-28376), rabbit anti-GluA1 (Millipore, AB1504), mouse anti-GluA2 (NeuroMab, 75-002), mouse anti-GluN1 (BD, 556308), mouse anti-GluN2A (NeuroMab, 75-288), rabbit anti-GluN2B (Millipore, 06-600), mouse anti-MAP2 (Abcam, ab11268), rabbit anti-mGlu5 (Millipore, AB5675), mouse anti-Nestin (Millipore, MAB5326), mouse anti-NLGN1 (NeuroMab, 75-160), mouse anti-Oct4 (Santa Cruz Biotechnology, sc-5279), mouse anti-Pin1 (Santa Cruz Biotechnology, sc-46660), mouse anti-PSD-95 (NeuroMab, 75-028), rabbit anti-Sox2 (Proteintech, 11064-1-AP), rabbit anti-SULT4A1 (Proteintech, 12578-1-AP), rabbit anti-VGAT (Synaptic System, 131 003).

All HRP- and fluorophore-conjugated secondary antibodies were purchased from Jackson ImmunoResearch Laboratories.

### Statistics

Data are expressed as Mean ± SEM or percentage, were analyzed for statistical significance, and were displayed by Prism 7 software (GraphPad, San Diego, CA). Shapiro-Wilk test or D’Agostino-Pearson test were applied to test the normal distribution of experimental data. Normal distributions were compared with *t*-test or ANOVA with appropriate *post-hoc* test. Non normal distributions were compared with the non-parametric Mann-Whitney test or Kruskal-Wallis test with appropriate *post-hoc* test, as indicated. The accepted level of significance was *P*≤0.05.

Statistical analyses for electrophysiological experiments were performed with OriginPro software. Distributions of cumulative probabilities were compared with Kolmogorov-Smirnov test.

## Acknowledgment

This work was supported by Comitato Telethon Fondazione Onlus; contract grant number: GGP16131 to C.V, Regione Lombardia; contract grant number:”AMANDA” CUP_B42F16000440005 to C.V; Regione Lombardia NeOn Progetto “NeOn” POR-FESR 2014-2020, “, ID 239047, CUP E47F17000000009 to C.V.

## Competing interests

No competing interests declared

## Supplementary Figures

**Figure 1 figure supplement 1.**
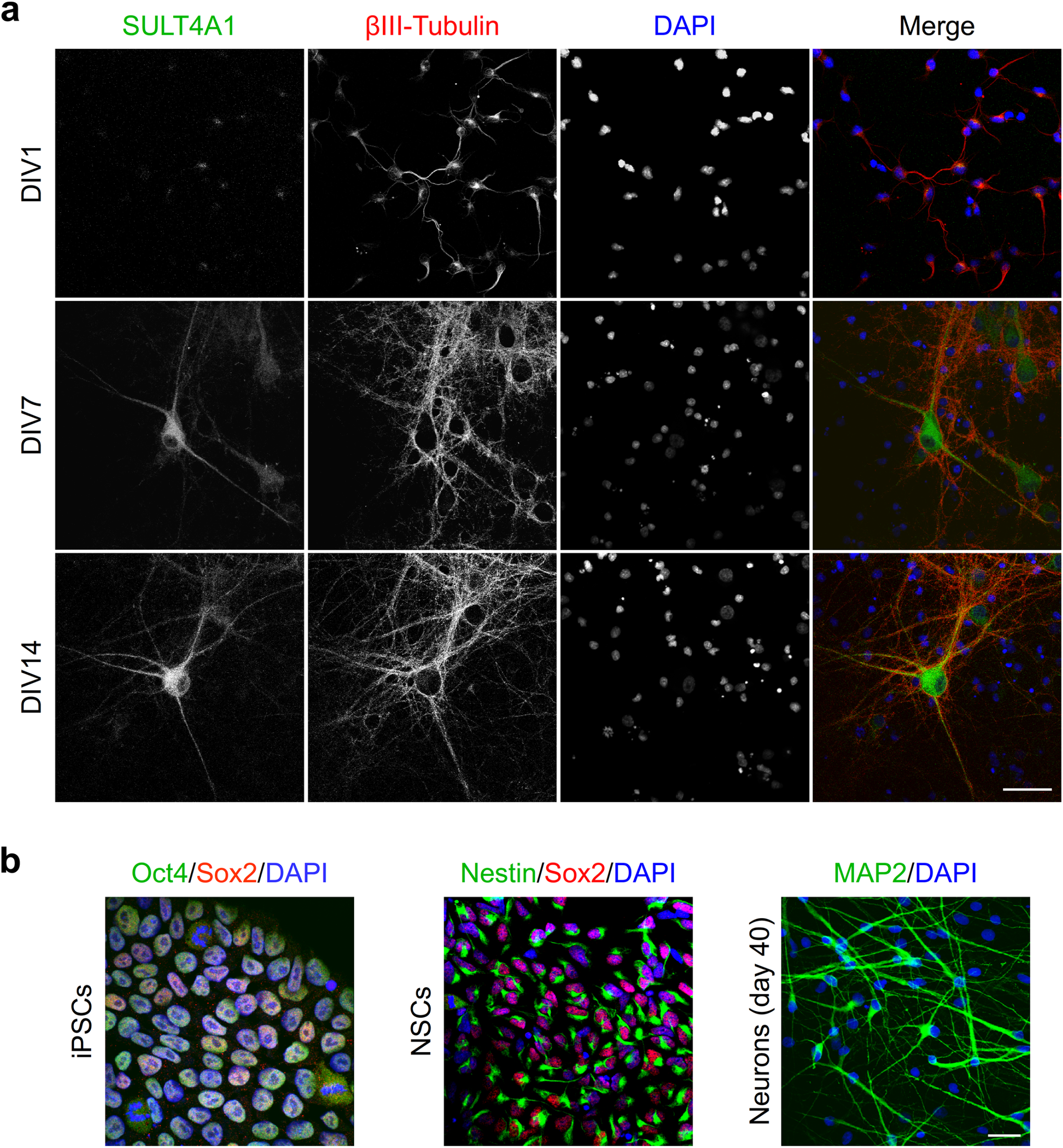
Physiological expression of SULT4A1 in primary cortical neurons and iPSC-derived neurons. (**a**) Representative immunocytochemical staining for SULT4A1 (green), βIII-Tubulin (red) and DAPI (blue), in rat cortical neurons at day-in-vitro (DIV) 7 and 14. (**b**) Left: representative immunocytochemical staining of Oct4 (green), Sox2 (red) and DAPI (blue) of induced pluripotent stem cells (iPSCs) obtained from healthy control individuals. Center: iPSC-derived neural stem cells (NSCs) stained for Nestin (green), Sox2 (red) and DAPI (blue). Right: representative immunostaining for MAP2 (green) and DAPI (blue) of NSC-derived neurons after 40 days of differentiation. Scale bar = 50μm.

**Figure 2 figure supplement 1.**
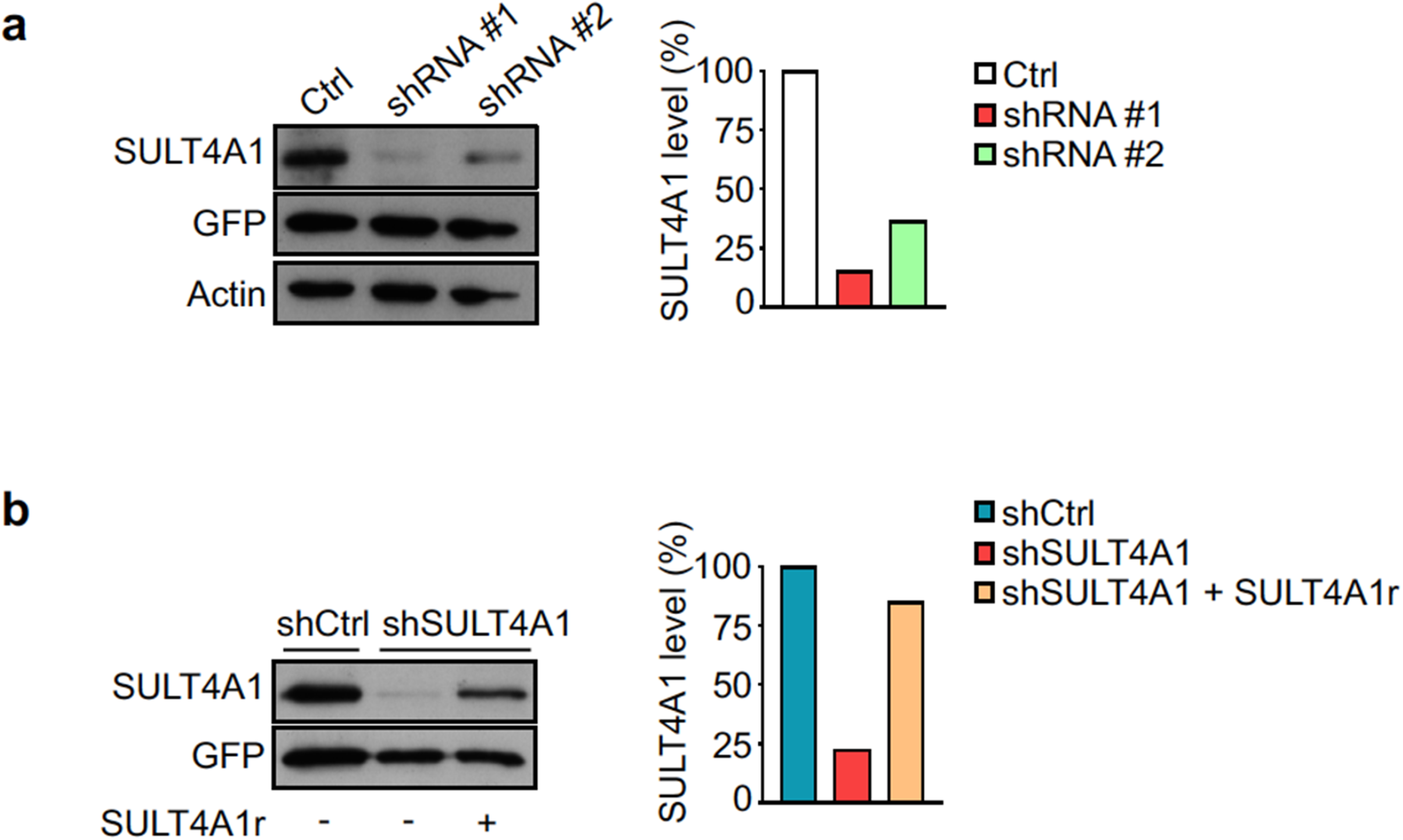
SULT4A1-specific shRNA validation. (**a**) Western blot from HEK293T cells transfected with a plasmid expressing pLVTHM (Ctrl) or one of the two shRNA specific for SULT4A1 (shRNA #1 or shRNA #2). All constructs express GFP. SULT4A1 protein level is graphed as percentage compared to the control condition (Ctrl). (**b**) Western blot from HEK293T cells transfected with a plasmid expressing the scrambled form of the shRNA (shCtrl), the shRNA #1 (now indicated as shSULT4A1) alone or together with the construct resistant to interference by shSULT4A1 (SULT4A1r). Both shCtrl and shSULT4A1 constructs also express GFP. SULT4A1 protein level is graphed as percentage compared to the control condition (shCtrl). Detailed statistical information for represented data is shown in Supplementary file 1

**Figure 2 figure supplement 2.**
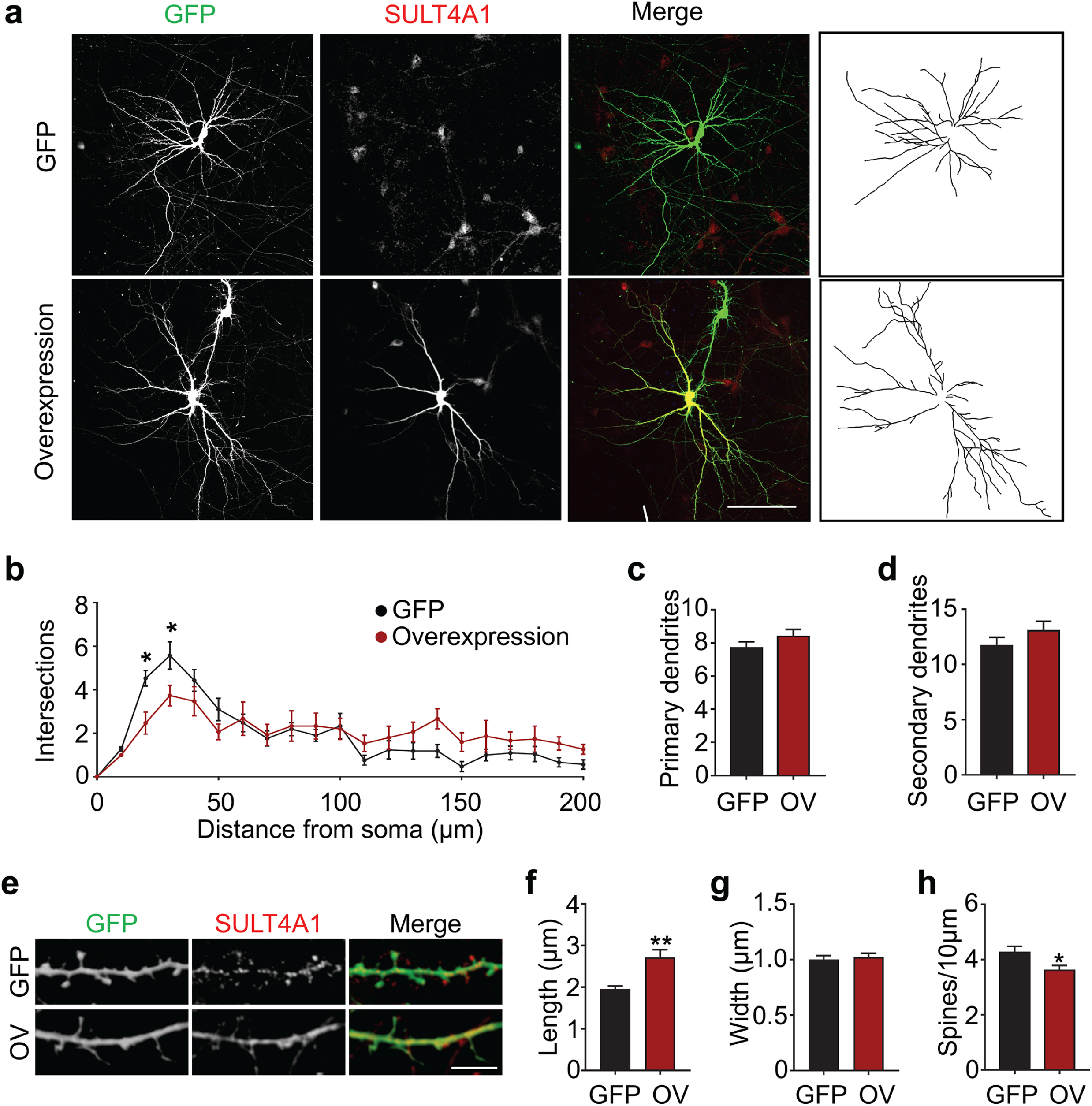
SULT4A1 overexpression impairs dendritic arborization and spines. (**a**) Representative immunofluorescence images and relative traces of rat cortical neurons transfected with pLVTHM alone (GFP) or together with pFlag-SULT4A1 (overexpression). Neurons were stained for SULT4A1 (red) to assess protein (over)expression. Scale bar= 100 μm. (**b**) Results of Sholl analysis and quantification of the number of intersections, plotted against the distance from the soma (GFP n=21, Overexpression n=15; Two-way ANOVA, *P*<0.0001; Sidak’s *post-hoc* test, * *P*<0.05). (**c-d**) Quantification of the number of primary (c) and secondary (d) dendrites (GFP n=20, Overexpression (OV) n=21; for (c) Mann-Whitney test, P=0.3511; for (d) Unpaired *t* test, *P*=0.2496). (**e**) Representative immunofluorescence images showing dendritic spines of rat cortical neurons transfected with pLVTHM alone (GFP) or together with pFlag-SULT4A1 (OV). Neurons were stained for SULT4A1 (red) to assess protein (over)expression. Scale bar= 10 μm. (**f-h**) Quantification of dendritic spines length (f), width (g) and number of spines per 10 μm of dendrite (h) (GFP n=15, OV n=23; Unpaired *t* test, * *P*<0.05, ** *P*<0.01). Data represent Mean ± SEM. Detailed statistical information for represented data is shown in Supplementary file 1

**Culotta et al. SULTA4A1 modulates synaptic development and function by promotir**

**Figure 2 figure supplement 2 - Source Data 1**

**Figure 2 figure supplement 2b.**
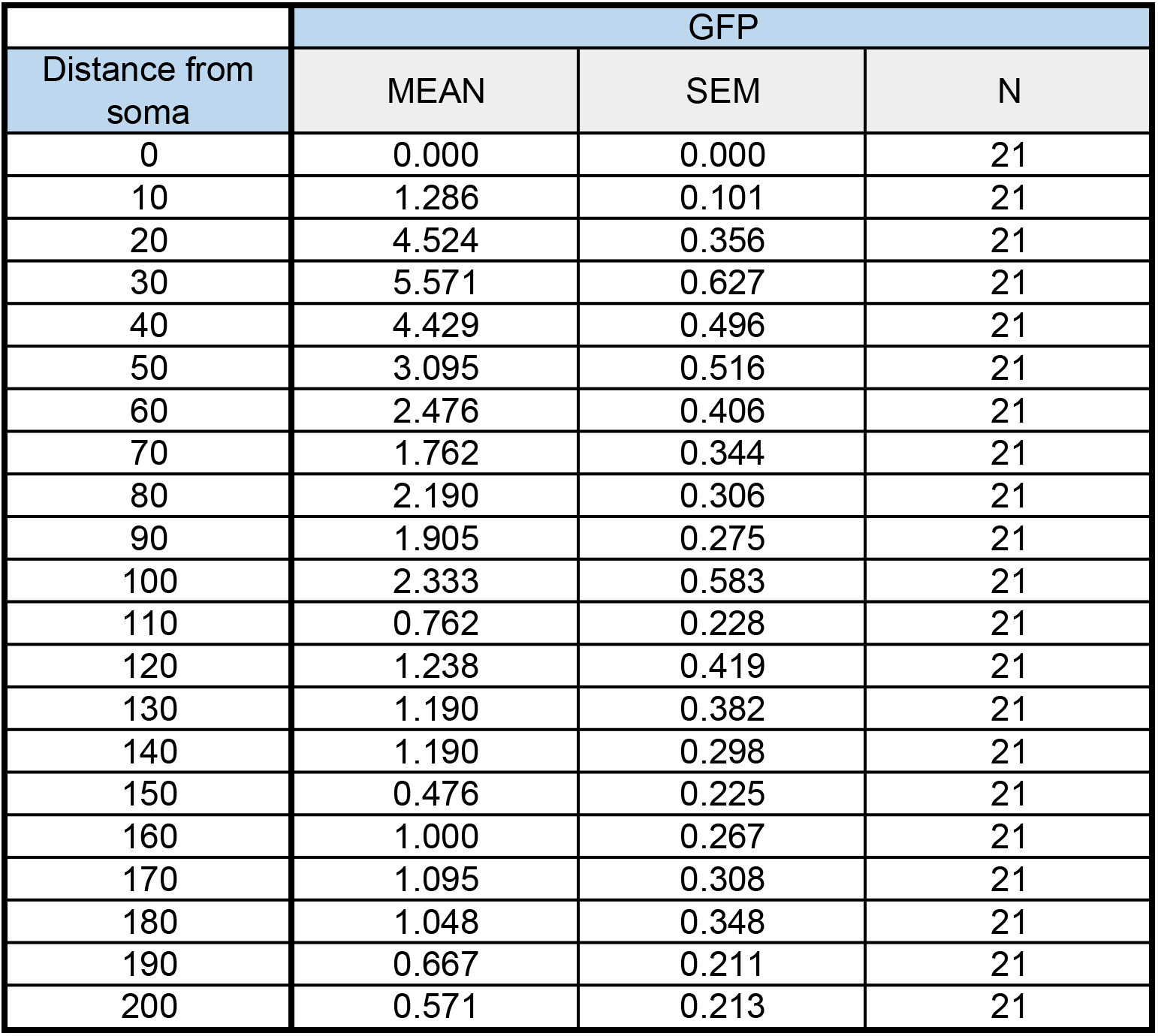
Number of intersections in rat cortical neurons

**Figure 2 figure supplement 2c.**
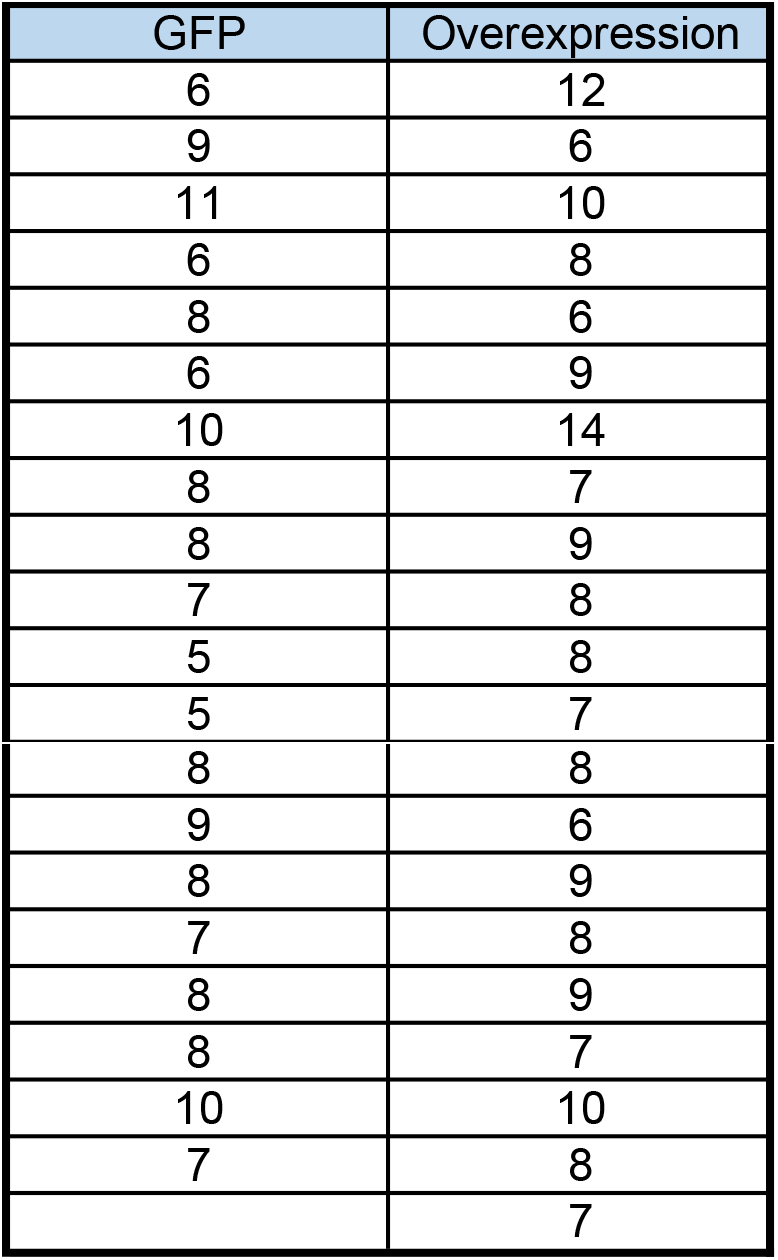
Number of primary dendrites in rat cortical neurons

**Figure 2 figure supplement 2d.**
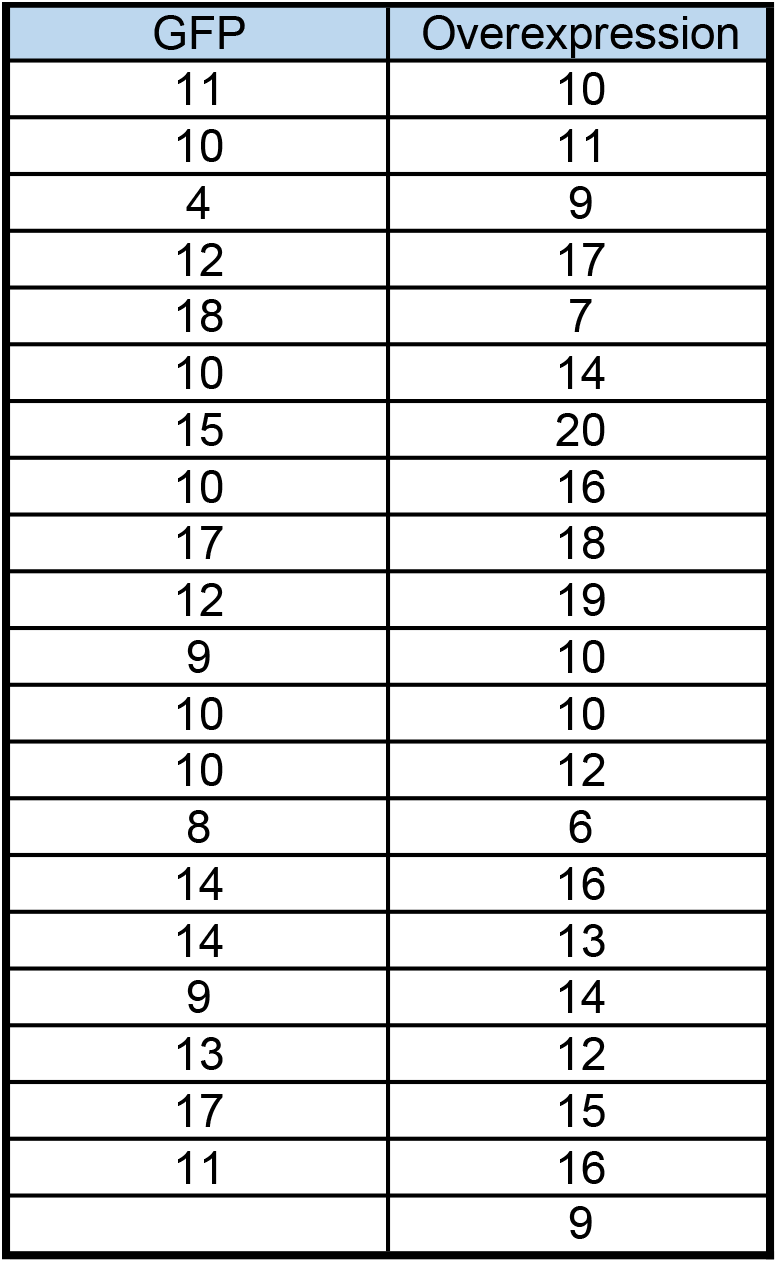
Number of secondary dendrites in rat cortical neurons

**Figure 2 figure supplement 2f.**
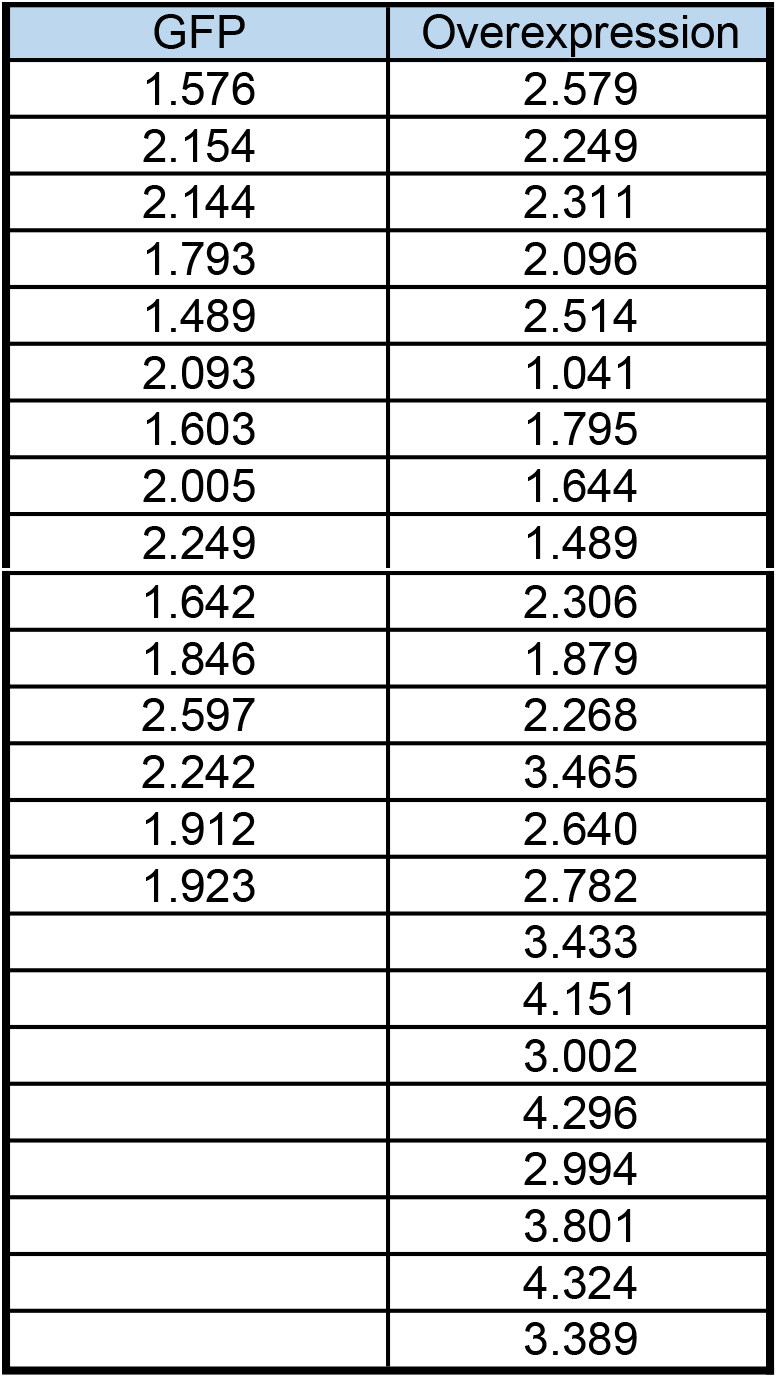
Dendritic spine length in rat cortical neurons (μm)

**Figure 2 figure supplement 2g.**
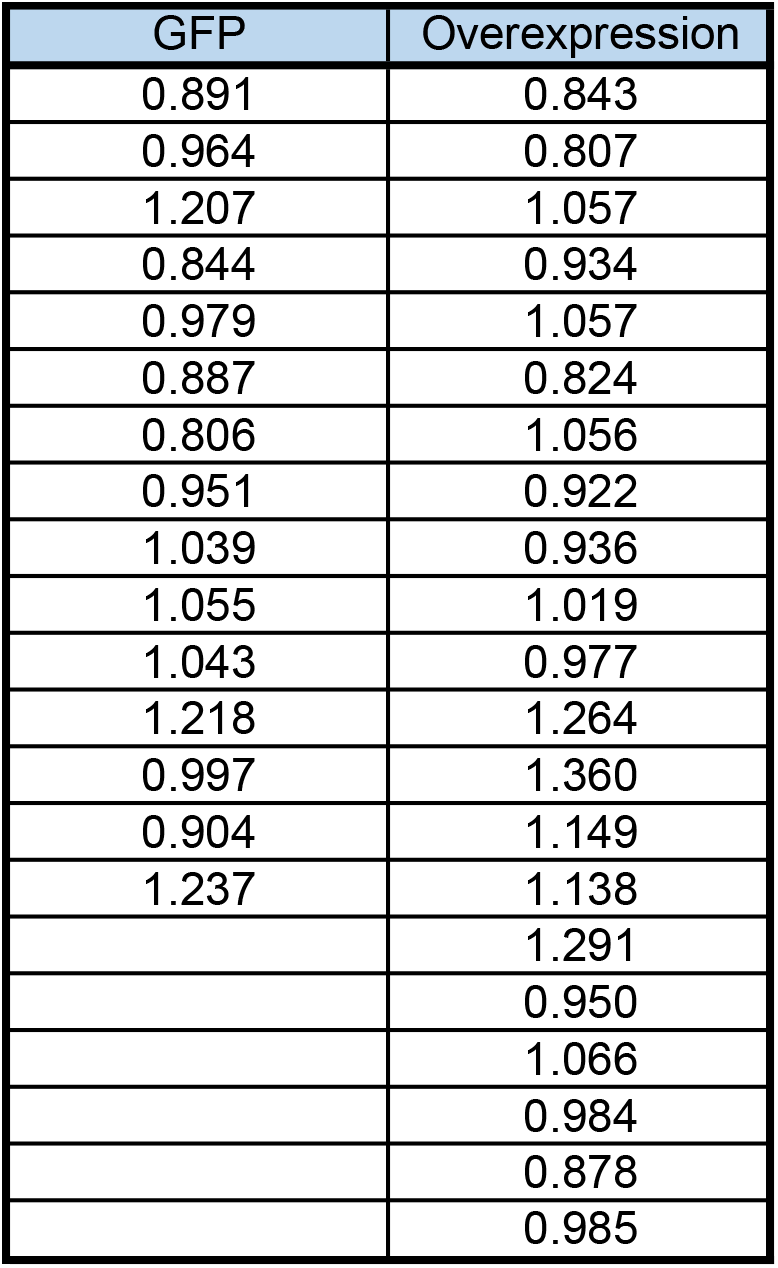
Dendritic spine width in rat cortical neurons (μm)

**Figure 2 figure supplement 2h.**
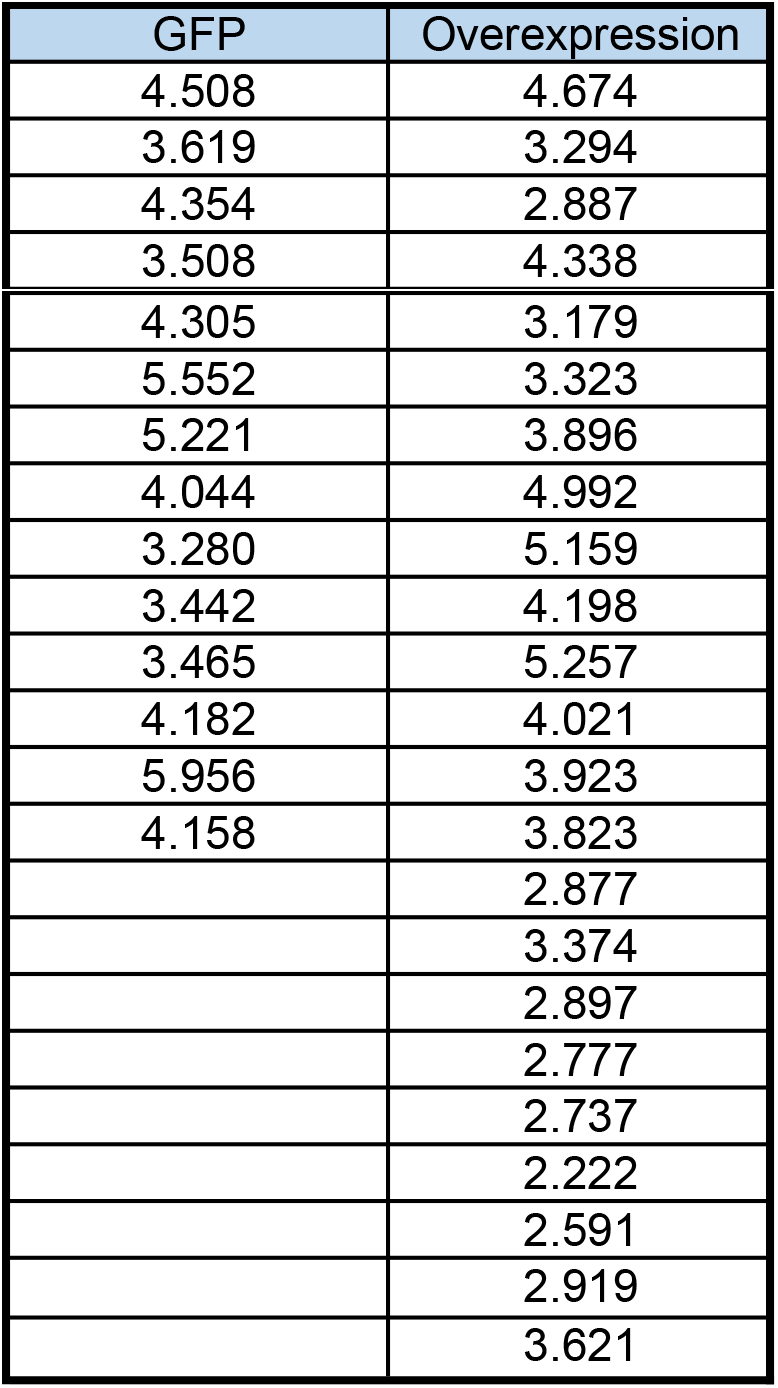
Dendritic spine density in rat cortical neurons (number of spines/10μm)

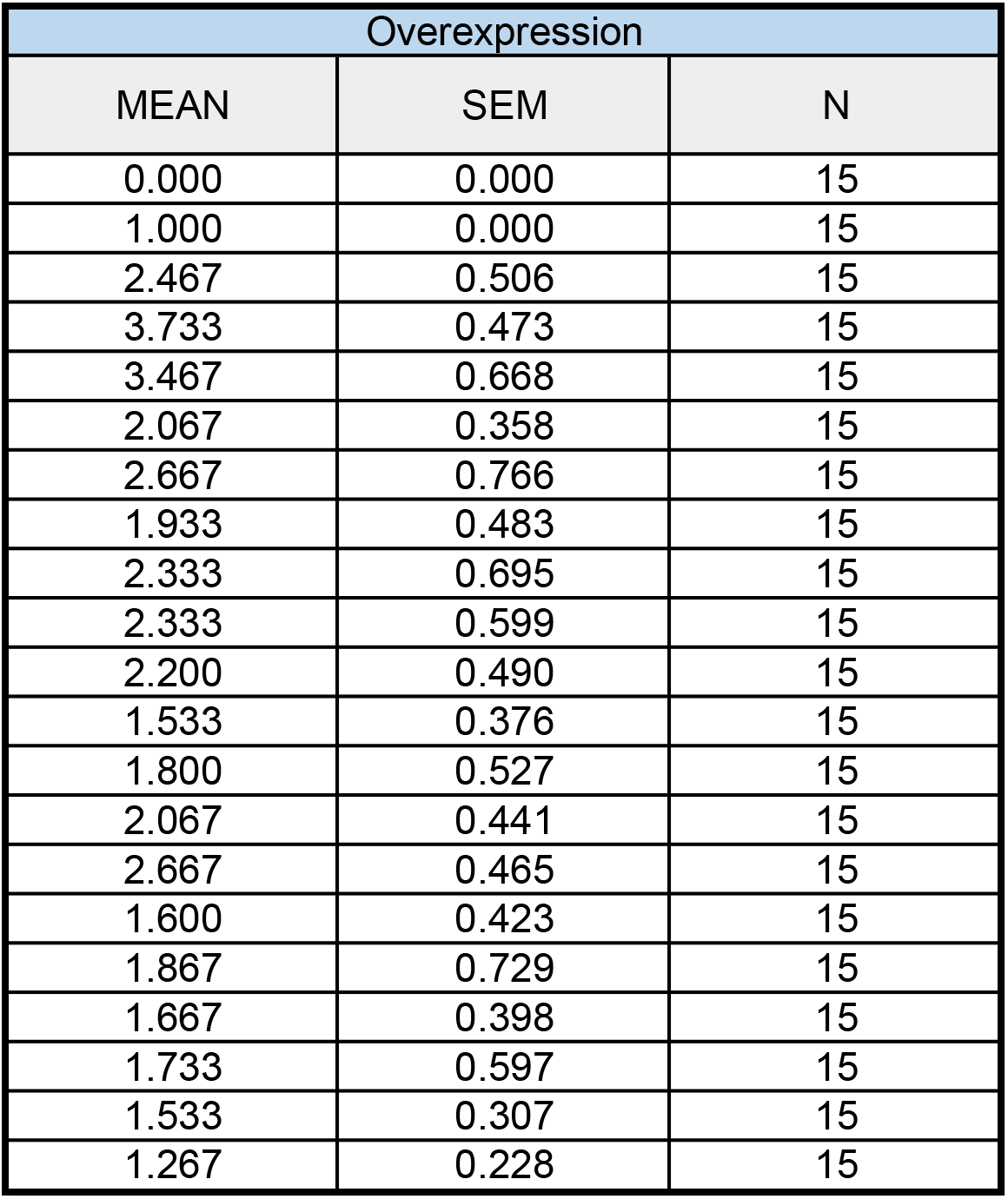

ng the formation of PSD-95/NMDAR complex

